# PCIF1 catalyzes m6Am mRNA methylation to regulate gene expression

**DOI:** 10.1101/484931

**Authors:** Erdem Sendinc, David Valle-Garcia, Abhinav Dhall, Hao Chen, Telmo Henriques, Wanqiang Sheng, Karen Adelman, Yang Shi

## Abstract

mRNA modifications play an important role in regulating gene expression. One of the most abundant mRNA modifications is N6,2-O-dimethyladenosine (m6Am). Here, we demonstrate that m6Am is an evolutionarily conserved mRNA modification mediated by the Phosphorylated CTD Interacting Factor 1 (PCIF1), which catalyzes m6A methylation on 2-O-methylated adenine located at the 5’ ends of mRNAs. Furthermore, PCIF1 catalyzes only 5’ m6Am methylation of capped mRNAs, but not internal m6A methylation *in vitro* and *in vivo*. Our global mRNA methylation analysis revealed that there is no crosstalk between m6Am and m6A mRNA methylation events, suggesting that m6Am is functionally distinct from m6A. Importantly, our data indicate that m6Am negatively impacts translation of methylated mRNAs by antagonizing cap binding protein eIF4E. Together, we identify the first and only human mRNA m6Am methyltransferase and demonstrate a novel mechanism of gene expression regulation through PCIF1-mediated m6Am mRNA methylation in eukaryotes.

**Highlights:**

- PCIF1 is an evolutionarily conserved mRNA m6Am methyltransferase
- Loss of PCIF1 leads to a complete loss of m6Am, whereas m6A level and distribution are not affected
- PCIF1 mediated m6Am does not affect RNA Pol II transcription or mRNA stability
- m6Am-Exo-Seq is a robust methodology that enables global m6Am mapping
- m6Am suppresses cap dependent translation

## Introduction

An important step in the regulation of gene expression is accomplished through chemical modifications of mRNAs. One of the best studied internal mRNA modifications is N6-methyladenosine (m6A), which is present on a diverse set of mRNAs, typically clustered around stop codons (Dominissini et al., 2012; Meyer et al., 2012). m6A is catalyzed by METTL3/METTL14 complex, and this modification regulates gene expression by influencing localization, stability, splicing or translation of mRNAs (Yang et al., 2018). Another abundant mRNA modification is N6,2-O-dimethyladenosine (m6Am), which occurs near the mRNA cap (Wei et al., 1975). 5’ ends of eukaryotic mRNAs typically carry a 7-methylguanosine (m7G) cap linked to the rest of the mRNA by a triphosphate linkage. The first nucleotide after the m7G cap can be methylated on the ribose sugar. If this first nucleotide is 2-O-methyladeonisine (Am), it can be further methylated at its N6 position to generate m6Am. While the methylase mediating Am methylation is known, the methyltransferase responsible for generating m6Am has not been identified (Keith et al., 1978; Mauer et al., 2017). As a consequence, the impact of m6Am on gene regulation is still largely unexplored.

Using multiple approaches, we provide both *in vitro* and *in vivo* evidence supporting Phosphorylated CTD Interacting Factor 1 (PCIF1) as the one and only mammalian m6Am methyltransferase. In order to investigate the distribution of m6Am, we developed a novel transcription-wide mapping technique we name m6Am-Exo-Seq. PRO-Seq, RNA-seq and m6Am-Exo-Seq analyses carried out in wildtype and PCIF1 null cells show that mRNA stability of m6Am-enriched genes is largely unaffected by the loss of PCIF1. In contrast, reporter assays carried out in PCIF1 null cells using mRNA transcripts with and without m6Am suggest that m6Am may function to suppress protein translation. Taken together, our findings identify a novel enzyme that mediates m6Am methylation in the nucleus, which impacts translation in the cytoplasm.

## Results

### m6Am is an evolutionarily conserved RNA modification

To study the biological significance of m6Am, we developed a highly sensitive mass spectrometry approach (LC-MS/MS) to detect and quantify m6Am (Figure S1). The triphosphate bond between the m7G cap and the first nucleotide of the mRNA is not cleaved by the enzymes typically used to generate single nucleosides for LC-MS/MS analysis. Therefore, without the removal of the m7G cap, only the internal m6A and Am modifications are detectable in human mRNA (Figure 1B). On the other hand, m6Am is detectable only after treatment with a de-capping enzyme, indicating that m6Am is restricted to the first nucleotide adjacent to the m7G cap of the mRNA (Figure 1B) (Linder et al., 2015). Using this protocol, we analyzed mRNA isolated from various model organisms by LC-MS/MS. While m6Am was not detected in fission yeast, nematode or Drosophila samples, mRNA isolated from zebrafish, mouse and human cells yielded m6Am, indicating that m6Am is an evolutionary conserved mRNA methylation (Figure 1C).

**Figure 1.**
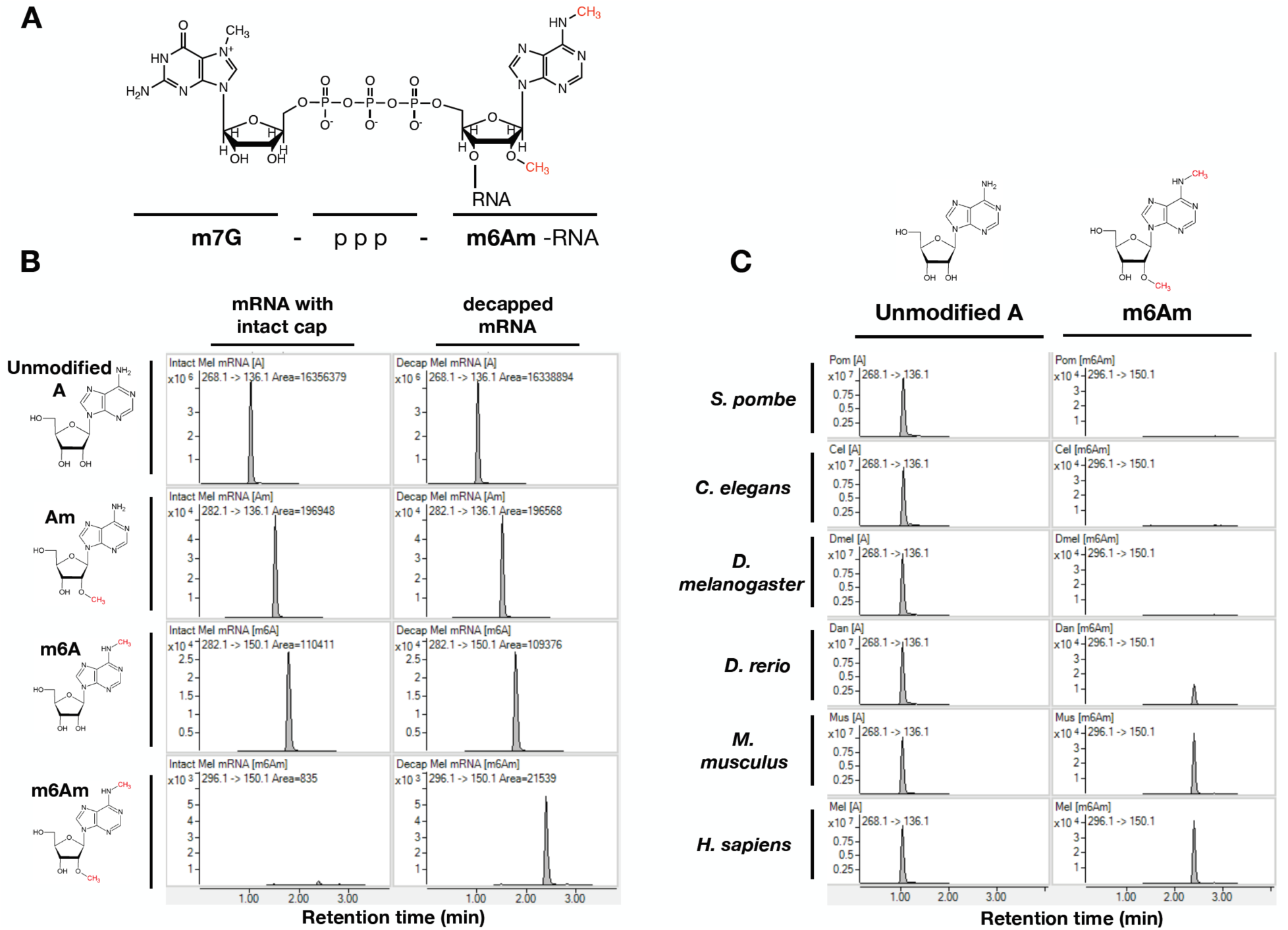
m6Am is an evolutionarily conserved RNA modification that is restricted to 5’ end of mRNAs. 1A. Chemical structure of 5’ end of eukaryotic mRNA with m6Am methylation. 1B. MS spectra of mRNA from MEL624 human melanoma cell line with or without treatment by de-capping enzyme to remove m7G cap prior to digestion for MS analysis. m6Am is detectable only when mRNA is de-capped. 1C. MS spectra of de-capped mRNAs isolated from indicated organisms.

### PCIF1 is required for mRNA m6Am methylation *in vivo*

A putative methyltransferase enzyme that is conserved in zebrafish, mouse and human, but absent in yeast and nematodes, is PCIF1 (Phosphorylated CTD Interacting Factor 1) (Figure S2) (Iyer et al., 2016). PCIF1 contains a N-terminal WW domain, which is believed to mediate interactions with RNA Polymerase II (Pol II) phospho-CTD, and also a putative C-terminal methyltransferase domain (Ebmeier et al., 2017; Fan et al., 2003).

To determine whether PCIF1 is a m6Am methyltransferase, we generated several independent PCIF1 knockout (KO) clonal cell lines in human melanoma MEL624 cells and determined its mRNA m6Am level by LC-MS/MS (Fig. 2A and 2B). mRNA isolated from a control MEL264 cell line yielded around 0.04% m6Am/A ratio. Strikingly, knocking out PCIF1 resulted in a complete loss of this modification in mRNA samples (Figure 2B). We further verified the absence of m6Am in PCIF1 KO cells by performing thin layer chromatography (TLC) (Figures 2C and S3A). Although PCIF1 KO resulted in a complete loss of m6Am, global m6A levels did not change, suggesting that PCIF1 is not required for m6A modification and that m6A is not dependent on m6Am levels (Figure S3B).

**Figure 2.**
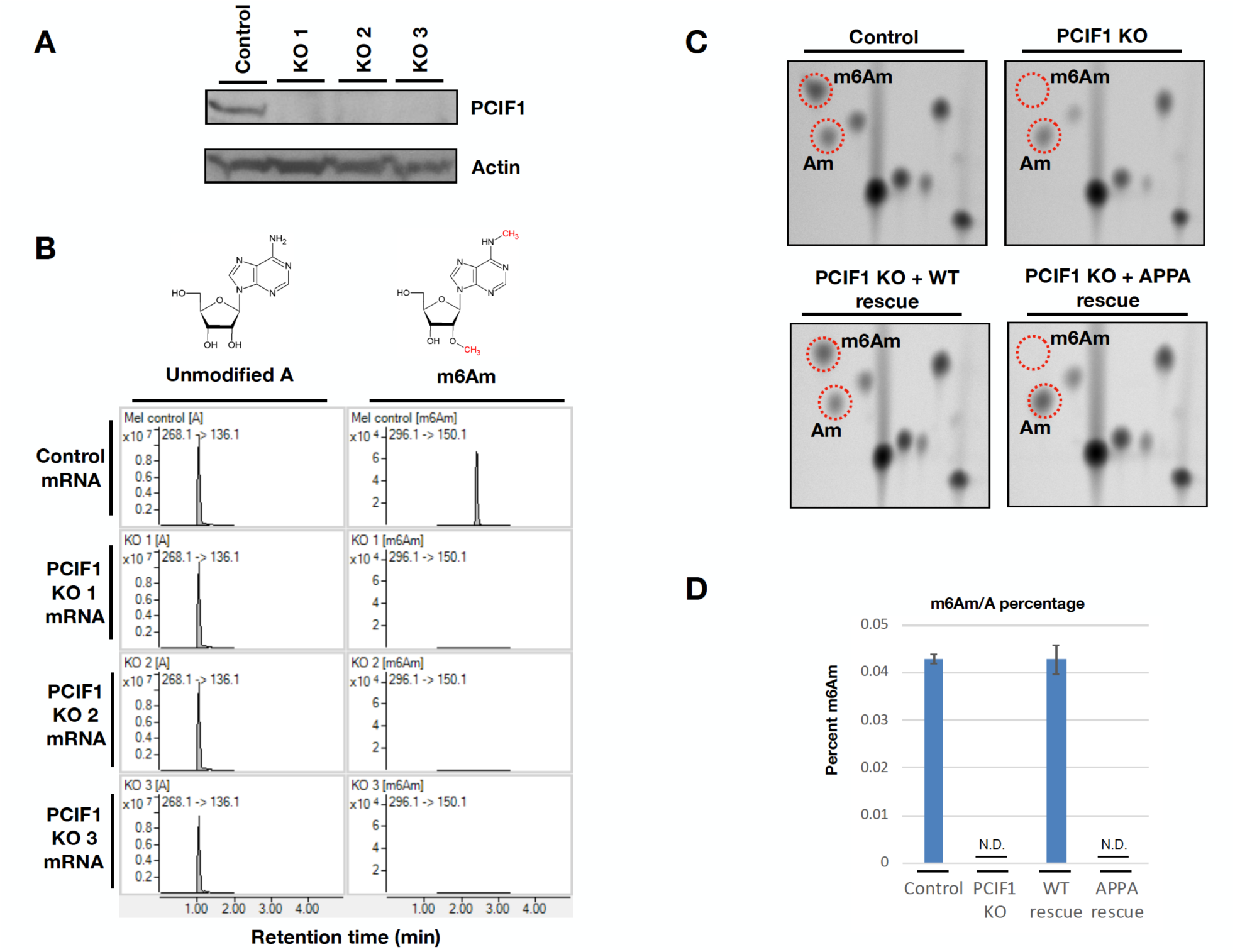
PCIF1 is required for mRNA m6Am methylation *in vivo*. 2A. Western blot analysis of control and three independent PCIF1 KO MEL624 cell lines with antibodies against endogenous PCIF1. Beta-actin is shown as a loading control. 2B. MS spectra of de-capped mRNA from control and three independent PCIF1 KO MEL624 cell lines. 2C. Thin-layer chromatography analysis of cap-adjacent nucleotides from mRNA isolated from control, PCIF1 KO and KO cells rescued with wild type or catalytic mutant PCIF1 (APPA). 2D. HPLC-MS/MS m6Am/A percentages of mRNAs from control, PCIF1 KO and KO cells rescued with wild type or catalytic mutant PCIF1 (APPA). (N.D., Not Detected). (n=3). Error bars depict standard error.

To determine whether it is the intrinsic enzymatic activity of PCIF1 that is responsible for m6Am methylation, we carried out rescue experiments by introducing either wild-type (WT) or catalytically inactive PCIF1 (APPA) back into the KO cells. While the expression of wild-type PCIF1 restored the global m6Am level in the KO cells, expression of the catalytic mutant with the conserved methyltransferase NPPF motif altered to APPA did not result in m6Am detectable by mass spectrometry or TLC (Figures 2C and 2D). PCIF1 contains a nuclear localization signal and has been suggested to be a nuclear protein (Ebmeier et al., 2017; Fan et al., 2003; Hirose et al., 2008). Consistently, we confirmed that PCIF1 is predominantly nuclear localized in both rescue cell lines by immunofluorescence (Figure S3C). These results indicate that PCIF1 is required for mRNA m6Am modification, which likely takes place in the nucleus *in vivo*.

### Recombinant PCIF1 methylates capped mRNA *in vitro*

To determine whether this putative methyltransferase can directly catalyze m6Am, we purified full length recombinant PCIF1 from bacteria (Figure S4A) and performed *in vitro* methylation assays employing firefly luciferase mRNA with m7G-Am 5’ cap as a substrate. Wild-type recombinant enzyme generated m6Am efficiently, whereas the APPA catalytic mutant did not show any activity (Figures 3A and S4B). Furthermore, PCIF1 did not generate m6A or Am modifications. On the other hand, generation of m6Am was accompanied by a decrease in the Am level, indicating that PCIF1 catalyzed conversion of Am to m6Am (Figure 3A). Consistent with previous findings (Kruse et al., 2011; Mauer et al., 2017), we failed to detect cap-adjacent m6A by TLC and PCIF1 KO cells show a greater accumulation of cap-adjacent Am (Figure S3A). These *in vitro* and *in vivo* data collectively suggest that PCIF1 specifically methylates Am-marked mRNAs.

**Figure 3.**
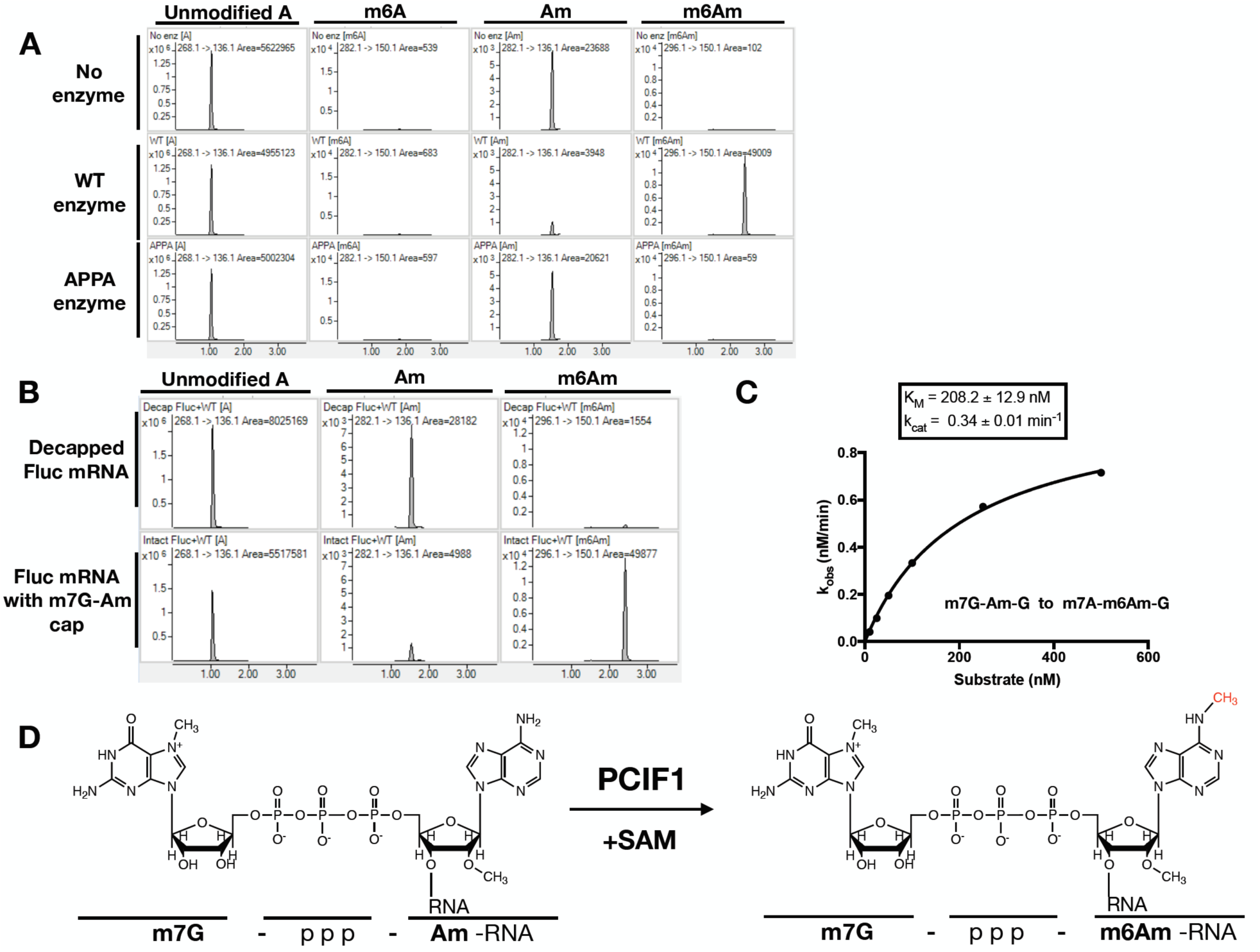
Recombinant PCIF1 methylates capped mRNA *in vitro*. 3A. Firefly luciferase mRNA with m7G-Am cap structure is methylated with wild-type or catalytic mutant recombinant PCIF1 *in vitro*. The methylation pattern of mRNA was analyzed by LC-MS/MS. 3B. In vitro methylation assays were performed with recombinant PCIF1 enzyme using de-capped firefly luciferase mRNA or mRNA with intact m7G-Am cap. 3C. Michaelis–Menten kinetics of recombinant PCIF1 was determined using m7G-Am capped dinucleotide RNA oligo by LC-MS/MS analysis. 3D. Chemical structures of PCIF1 substrate and product.

We next investigated whether the m7G cap structure is required for the catalytic activity of PCIF1, using de-capped luciferase mRNA with 5’ Am in *in vitro* methylation assays. The Am residue next to the intact m7G cap was readily converted to m6Am by recombinant PCIF1. In contrast, the Am residue on de-capped mRNA was not converted to m6Am, indicating that m7G cap is required for PCIF1 catalytic activity (Figure 3B). To determine whether there is a size requirement for capped mRNA to serve as a substrate, we used a capped RNA dinucleotide (m7G-AmG) in *in vitro* methylation assays. PCIF1 efficiently generated m6Am with the dinucleotide as a substrate with a K_m_ of 208 nM and K_cat_ at 0.34 min^-1^ (Figure 3C). This indicates that capped transcripts as short as 2 nucleotides can be efficiently methylated by PCIF1 (Figure 3D). Given that recombinant PCIF1 efficiently methylates capped mRNAs *in vitro* and that deletion of PCIF1 results in a complete loss of mRNA m6Am methylation *in vivo*, we conclude that PCIF1 is the only mRNA m6Am methyltransferase in human cells.

### Loss of PCIF1 does not affect global mRNA m6A distribution *in vivo*

In order to identify the mRNAs that carry m6A and m6Am, we performed MeRIP-Seq with an antibody against m6A in control and PCIF1 KO cell lines. Given that m6A and m6Am are structurally similar, antibodies against m6A are expected to enrich for m6Am at the 5’ end of the mRNAs, as has been previously suggested (Linder et al., 2015). However, although we were able to identify internal m6A loci enriched around stop codons, as reported previously for other cell lines (Dominissini et al., 2012; Meyer et al., 2012), we observed a very modest enrichment at the 5’ end of mRNAs that was mostly maintained in the PCIF1 KO cells (Figures 4 and 5A). Therefore, it appears that the regular m6A MeRIP-Seq protocol does not efficiently capture m6Am-enriched transcripts. In agreement with our *in vitro* and *in vivo* results (Figures 3A and S3B), global m6A distribution did not change in two independent PCIF1 KO lines, compared to control cells (Figure 4). Deletion of PCIF1, and hence complete loss of m6Am, altered neither global m6A level (Figure S3B) nor genome-wide m6A distribution (Figure 4A), suggesting that m6A and m6Am are independent mRNA modifications.

**Figure 5.**
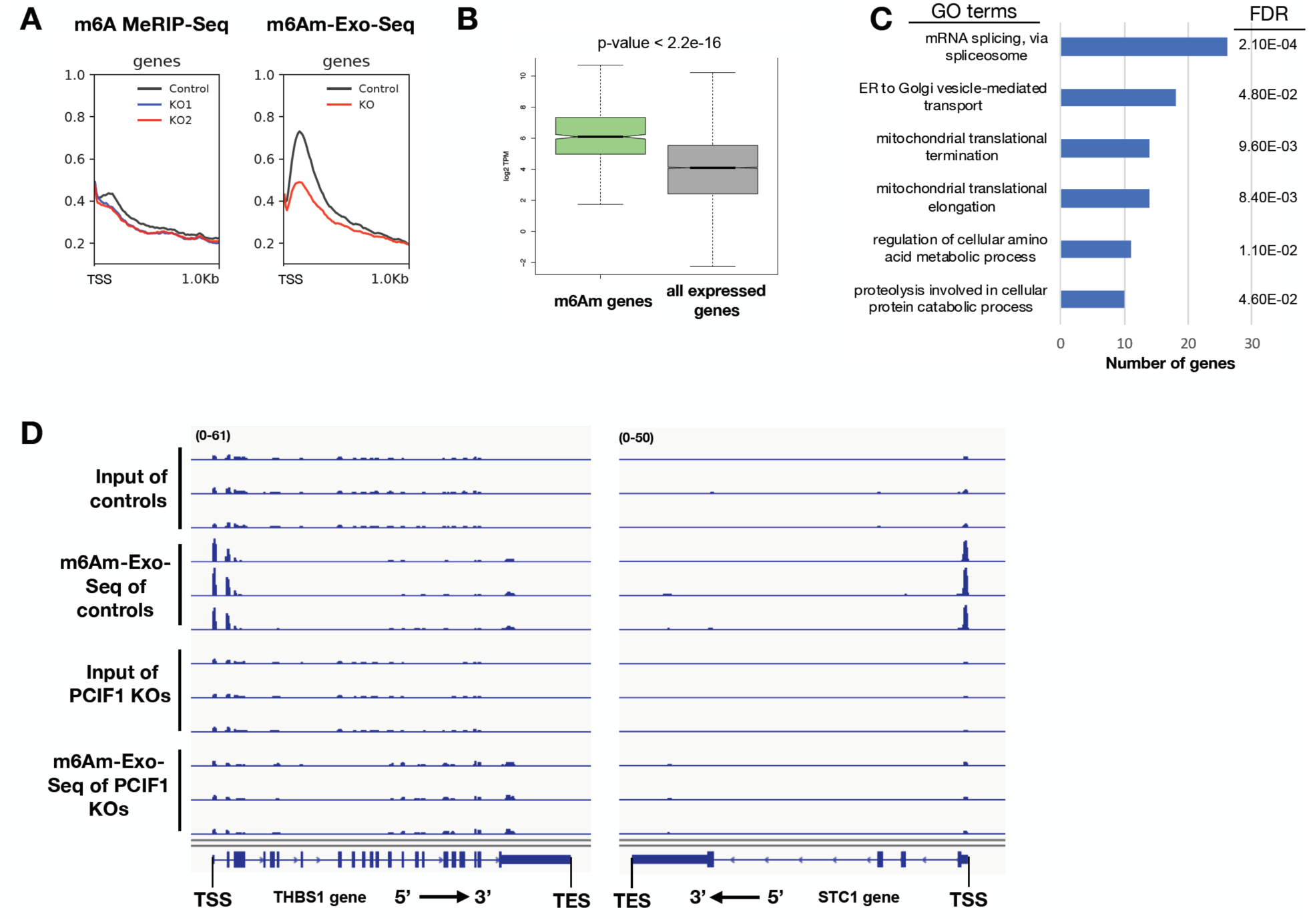
m6Am-Exo-Seq as a robust methodology that enables global m6Am mapping. 5A. Metagene plots of m6A(m) enrichment over Input near TSS of all expressed genes (n=15,258) in MEL624 cells employing either m6A MeRIP-Seq or m6Am-Exo-Seq methodology in control and PCIF1 KO cells. 5B. Boxplots of expression level (log2 TPM) of m6Am mRNAs (n=640) vs. all mRNAs expressed in MEL624 cell line (n=15,258). The observed difference is significant (p-value < 2.2e-16) using a Mann-Whitney test. Box plots show the 25th–75th percentiles and error bars depict the 10th–90th percentiles. 5C. Biological process Gene Ontology (GO) terms that have FDR<0.05 identified by DAVID 6.8 5D. Genome browser views of 2 example genes with m6Am enrichment using m6Am-Exo-Seq

**Figure 4.**
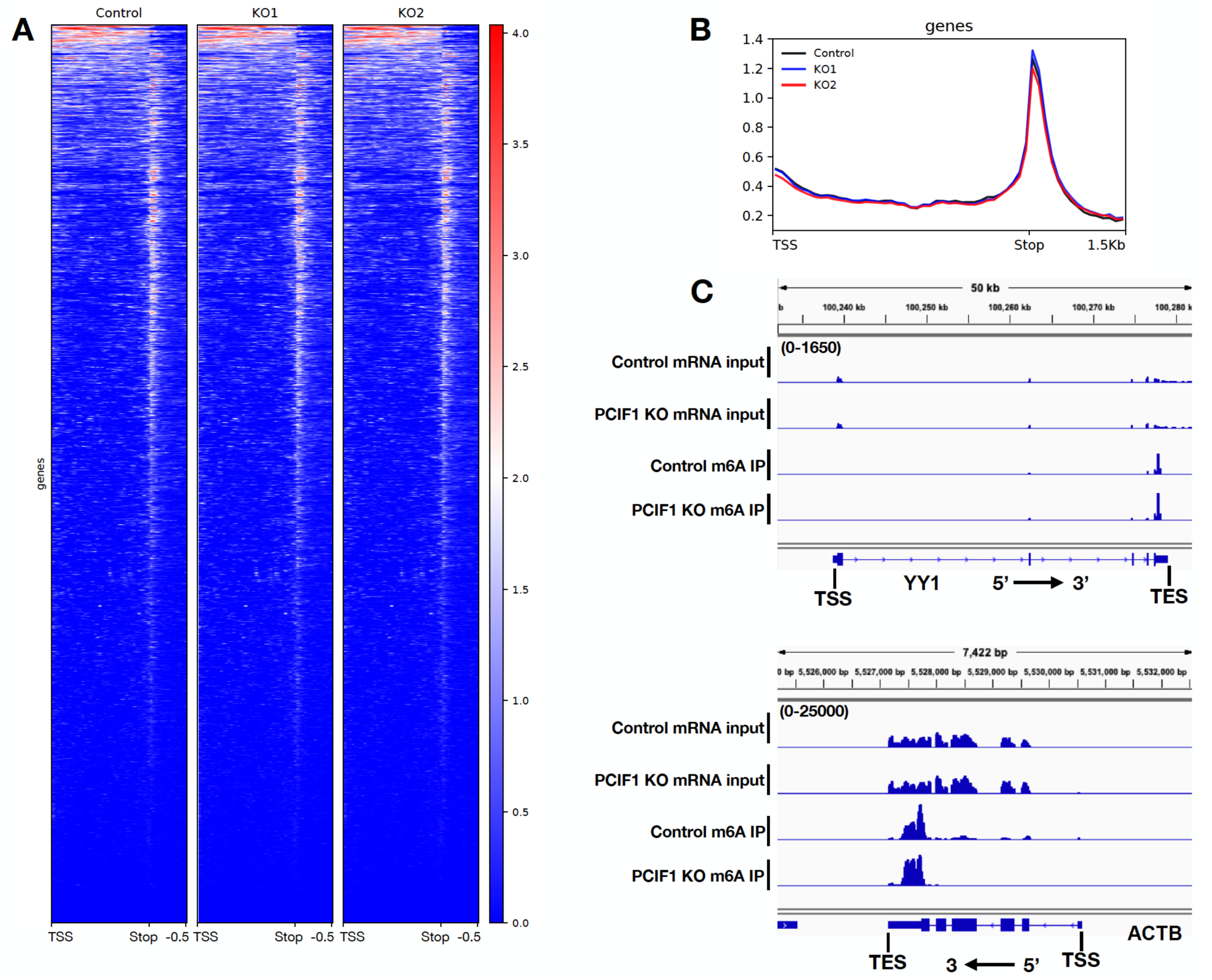
Loss of PCIF1 does not affect global mRNA m6A distribution *in vivo*. 4A. Heat map of m6A enrichment over Input of genes overlapping with m6A peaks (n=4,460) in MEL624 cell line determined by m6A MeRIP-Seq in control and two independent PCIF1 KO cell lines. 4B. Metagene plots of m6A enrichment over Input of genes overlapping with m6A peaks (n=4,460) in MEL624 cell line determined by m6A MeRIP-Seq in control and two independent PCIF1 KO cell lines. 4C. Genome browser views of two examples genes with m6A enrichment near stop codon.

### Development of m6Am-Exo-Seq as a robust methodology that enables global m6Am mapping

To be able to map m6Am modification transcriptome-wide, we developed a novel exonuclease assisted high-throughput sequencing methodology that we call m6Am-Exo-Seq (Figure S5). We performed m6Am-Exo-Seq on mRNA from control and PCIF1 KO cells using 8 different m6A antibodies. Our new technique allowed us to observe a clear enrichment of 5’ ends immediate to transcription start sites only in mRNAs from control cells, but not in mRNAs from PCIF1 KO cells, suggesting that m6Am-Exo-Seq specifically pull-down m6Am-enriched transcripts (Figure 5A). Seven out of the eight m6A antibodies tested with m6Am-Exo-Seq resulted in a significant enrichment of m6Am only in control mRNA, demonstrating the robustness of this novel methodology (Figure S5).

To globally map m6Am, we applied m6Am-Exo-Seq with a monoclonal m6A antibody in MEL624 cell line using three biological replicates of control and PCIF1 KO cells (Figure 5). We identified 640 genes that are enriched for this modification (Table S1). These transcripts are among the highly expressed genes in MEL624 cell line (Figure 5B), and are in diverse pathways including splicing, vesicular transport and mitochondrial metabolism (Figures 5B and 5C). m6Am was also mapped in HEK293T cells previously (Linder et al., 2015). While most of the m6Am transcripts are methylated in both HEK293T and MEL624 cells, around 40% of m6Am transcripts are unique to MEL624 cells (Figure S6A), suggesting that m6Am is a dynamic epitranscriptomic mark with cell-type specific patterns.

### PCIF1-mediated deposition of m6Am does not alter transcription or stability of target genes

PCIF1 interacts with the phosphorylated C-terminal domain (CTD) of Pol II, and has been implicated in negative regulation of gene expression (Ebmeier et al., 2017; Fan et al., 2003; Hirose et al., 2008). However, recent studies suggest that deposition of m6Am stabilizes mRNA or stimulates translation (Mauer et al., 2017), both of which would increase the expression of target genes. On the other hand, a separate study failed to find a correlation between m6Am methylation and mRNA stability (Wei et al., 2018). We thus wished to investigate the mechanism of m6Am activity in our system, where we can precisely abolish m6Am by PCIF1 KO, without perturbing m6A levels. We first performed RNA-Seq in PCIF1 KO MEL624 cells and a control cell line. We defined 236 mRNA species that were significantly more abundant in both PCIF1 KO cell lines, and 304 mRNAs with reduced abundance (Figure 6A, Tables S2 and S3). To distinguish whether these changes in mRNA levels were due to altered transcription mediated by m6Am, we next performed PRO-Seq in control and PCIF1 KO cells. PRO-Seq specifically measures engaged Pol II across the genome, providing definitive information on levels of active transcription (Kwak et al., 2013). These assays were spike normalized to enable accurate quantification and comparisons between samples. PRO-Seq signals across the 236 up-regulated genes from RNA-seq revealed that these genes were more highly transcribed in PCIF1 KO cells (Figure 6B). Likewise, down-regulated genes were less actively transcribed (Figure 6B). Importantly, there was a strong correlation between the changes observed in RNA-Seq and PRO-Seq, indicating that changes in RNA abundance could be explained fully by altered transcription (r=0.67, Figure 6D).

**Figure 6.**
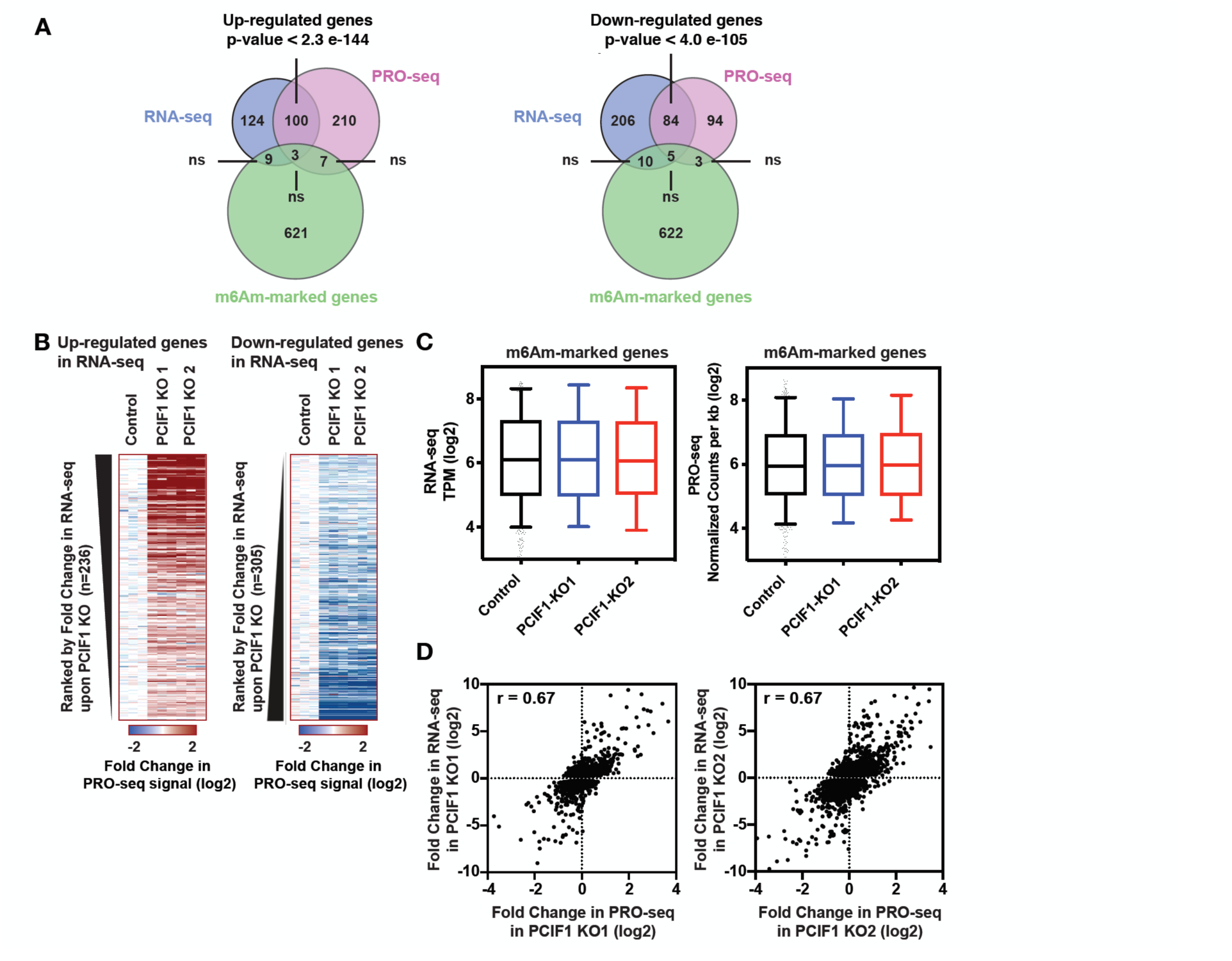
PCIF1-mediated deposition of m6Am does not alter transcription or stability of target genes. 6A. Venn diagrams displaying overlap between m6Am-enriched genes in MEL624 cells and genes identified as significantly upregulated (left) or downregulated (right) in both PCIF1 KO MEL624 cell lines, by RNA-Seq and PRO-Seq. P-values are calculated with a hypergeometric test. 6B. Heatmaps of PRO-Seq signal at genes found to be up- or down-regulated using RNA-Seq in PCIF1 KO cells. Shown are the fold changes in PRO-Seq signal across the bodies of genes up-regulated in RNA-Seq (left) or down-regulated in RNA-Seq (right). Genes are ranked from most upregulated to least (at left) and least down-regulated to most (at right), demonstrating that PRO-Seq signal scales with results from RNA-seq. 6C. RNA-Seq signal (left) and normalized PRO-Seq signal (right) within 640 m6Am-enriched genes in Control MELF624 cells, as compared to PCIF1 KO cells. Box plots show the 25th–75th percentiles and error bars depict the 10th–90th percentiles. Differences are not significant (P > 0.01). P-values were calculated using the Mann-Whitney test. 6D. Scatter plots displaying the correlation of RNA-Seq vs PRO-Seq KO/control fold change signal. Only genes that have a fold change > 1.2 according to RNA-Seq in KO1 or KO2 were plotted. The total number of genes used in KO1 plot is 5,774 and KO2 is 5,580. Both KOs have the same Pearson correlation value of 0.67.

Furthermore, we noted no significant overlap between m6Am-enriched genes and genes with significantly altered signals in RNA-Seq or PRO-Seq assays (Figure 6A, Tables S1-S5). PCIF1 KO did not change RNA-seq or PRO-Seq signal within m6Am-enriched genes as compared to controls (Figure 6C). Moreover, the loss of PCIF1 did not alter Pol II levels or distribution at noncoding snRNA genes that are marked by m6Am (Figure S6B) (Mauer et al., 2018; Wei et al., 2018). Taken together, our data suggest that although PCIF1 has been reported to bind RNA Pol II, it does not play a significant role in transcription at m6Am target genes. Moreover, steady-state transcript levels of m6Am-marked genes, as determined by RNA-Seq in PCIF1 KO cells, are not affected by the loss of this modification, suggesting that stability of m6Am-marked mRNAs is not significantly affected by PCIF1-mediated methylation.

### m6Am suppresses cap-dependent translation

Regulation of cap binding protein eIF4E during translation initiation is a key event in protein translation control. eIF4E binds m7G cap of mRNAs and recruits other translation initiation factors and ribosomes for translation initiation (Sonenberg and Gingras, 1998). The C terminal loop of cap bound eIF4E is in close proximity to the first nucleotide of the mRNA (Tomoo et al., 2002, 2005). The N6 residue of the cap-adjacent adenine of mRNA forms a critical hydrogen bond with threonine 205 residue of human eIF4E (Figure S7) (Tomoo et al., 2002, 2005). We thus hypothesized that modification of this adenine by PCIF1 to form m6Am may affect this interaction and hence impact translation at the initiation step. To test this, we used well established reporter assays to analyze the effect of m6Am on translation efficiency. We first transfected PCIF1 KO cells with EGFP mRNA with either m7G-Am or m7G-m6Am 5’ ends and determined the GFP signal by microscopy and flow cytometry (Figure 7A and S8). Although both mRNAs are transfected with similar efficiencies, m6Am-EGFP transfected cells showed diminished GFP signal (Figures 7A-D and S8), suggesting that m6Am-containing transcripts are translated less efficiently that the unmethylated counterparts *in vivo*. To further validate the effect of m6Am on translation and to investigate whether this effect is independent of the mRNA sequence, we performed *in vitro* translation assays using varying amounts of firefly luciferase mRNAs with cap adjacent Am or m6Am modifications, followed by luminometric quantification. Similar to our *in vivo* reporter assays, m6Am suppressed translation of luciferase mRNA *in vitro* (Figure 7E). Next, we investigated whether m6Am-mediated suppression is specific to cap-dependent translation by employing Am or m6Am methylated bi-cistronic mRNA transcripts in *in vitro* translation assays. m6Am suppressed the translation of cap-dependent firefly luciferase whereas IRES mediated translation of renilla luciferase was unaffected, indicating that m6Am specifically suppresses cap-dependent translation (Figure 7F).

**Figure 7.**
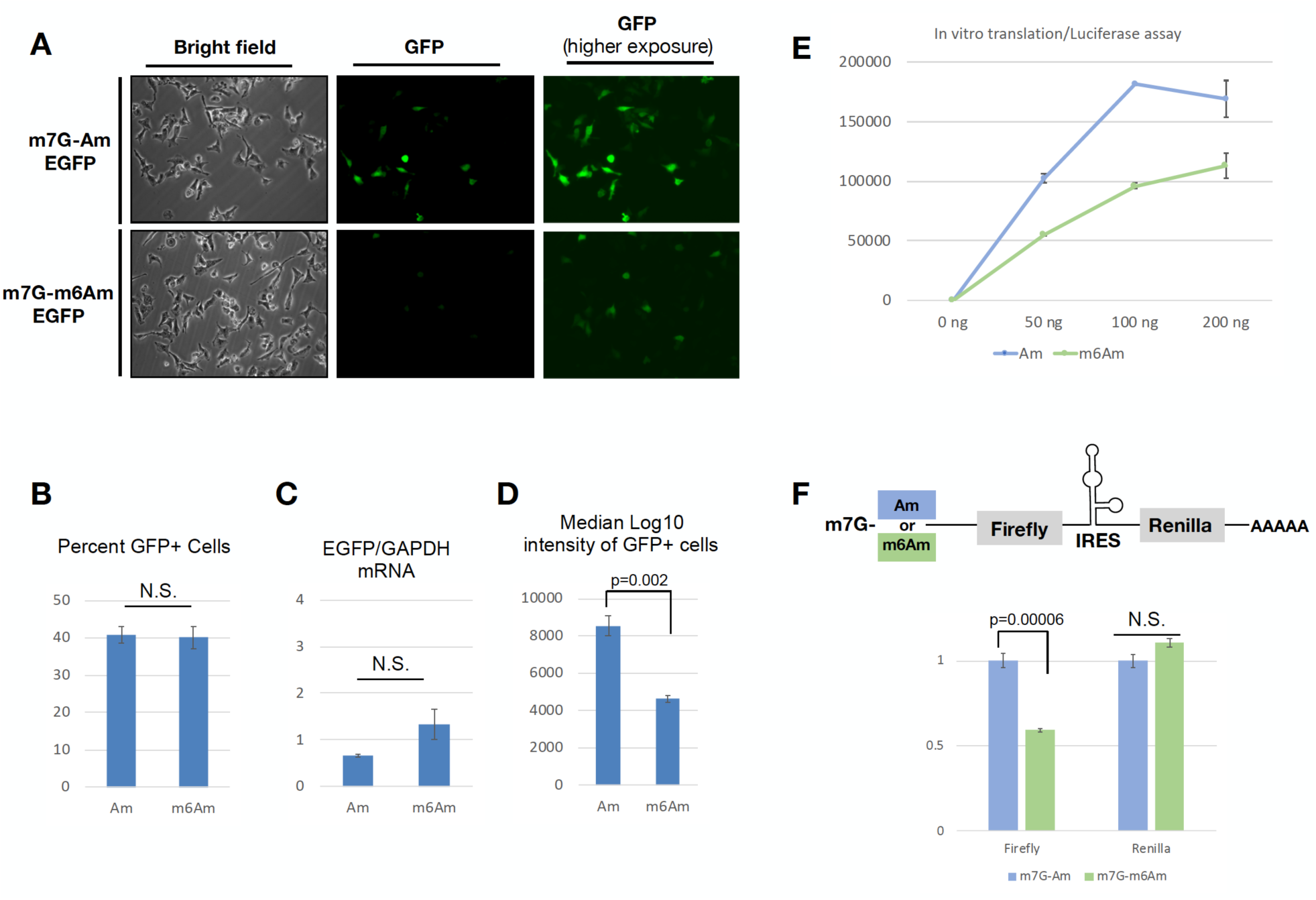
m6Am suppresses cap-dependent translation. 7A. EGFP mRNA that starts with either m7G-Am or m7G-m6Am was transfected into PCIF1 KO MEL624 cells. GFP signal was determined by fluorescence microscopy 24 hours post-transfection. 7B. The percentage of GFP positive cells from the experiment in 7A determined by flow cytometry (n=3 replicates). 7C. EGFP mRNA level from the cells in experiment 7A as determined by RT-qPCR. (n=3 replicates). 7D. Median fluorescence intensity of the cells from the experiment in 7A determined by flow cytometry. P-values in 7B-D are calculated using unpaired t-tests. (n=3 replicates). 7E. *In vitro* translation assays were carried out with indicated amounts of luciferase mRNA starting with m7G-Am or m7G-m6Am using rabbit reticulocyte system. Resulting luciferase signal was quantified on a plate reader. (n=4 replicates) 7F. In vitro translation assays were carried out with bicistronic construct that starts with m7G-Am or m7G-m6Am using rabbit reticulocyte system. Resulting firefly and renilla luciferase signals were quantified on a plate reader. (n=4 replicates). The p-values were calculated with an unpaired t-test. All error bars in this figure depict standard error.

Given that m6Am is a cap proximal RNA modification that suppresses cap-dependent translation we next determined whether m6Am affects the interaction between mRNA and cap binding protein eIF4E. Specifically, we analyzed binding dynamics between recombinant human eIF4E protein and capped Am and m6Am RNA oligos, as well as de-capped RNA oligos, by microscale thermophoresis (MST). As expected, the affinity of eIF4E to the RNA decreased significantly when the mRNA cap was removed. Importantly, we found that m6Am also decreased the affinity of eIF4E to the capped RNA oligo, in comparison to capped Am RNA (Figure S9). Taken together, these data suggest that m6Am modification suppresses cap-dependent mRNA translation, possibly by decreasing eIF4E binding.

## Discussion

In this study, we uncovered a new regulatory mechanism mediated by PCIF1 m6Am deposition on mRNAs. Our *in vivo* and *in vitro* data demonstrate that PCIF1 is the only methyltransferase necessary and sufficient for the conversion of cap-Am mRNAs to cap-m6Am mRNAs in human cells. Moreover, we provide evidence that m6Am negatively regulates cap dependent translation by decreasing the binding of eIF4E to m6Am-containing transcripts. Our findings highlight a previously unknown translational regulatory pathway that begins with a modification in the nucleus, opening new avenues of investigation in the epitranscriptome and translation fields.

Given that m6Am marks diverse transcripts including noncoding snRNAs and histone mRNAs that are translated independently of eIF4E, we predict that this modification will play diverse functions, which include but are not limited to protein translation. m6Am has been suggested to play a role in regulating the stability of modified mRNAs in HEK293T cells (Mauer et al., 2017). However, our analyses show that neither transcription nor abundance of m6Am-enriched transcripts are affected in PCIF1 KO MEL624 cells (Figure 6). However, we cannot rule out a potential role of m6Am in regulating mRNA stability under conditions that were not tested in this study.

Rather than altering mRNA stability, we found that m6Am suppresses cap dependent translation, possibly by reducing binding of eIF4E to the capped mRNAs. Our findings strongly suggest a role for PCIF1 in protein translation control, which will be validated by investigation of translation of endogenous mRNAs by PCIF1. Collectively, the identification of PCIF1 as the sole, evolutionarily conserved, mRNA m6Am methyltransferase paves the way for investigating the impact of PCIF1 on gene expression of m6Am-marked transcripts in diverse biological contexts.

## Acknowledgments

We thank members of the Shi lab for helpful discussions. We are especially indebted to the Tufts Genomic Core and Karla F Meza-Sosa for helpful advice and reagents. We thank Synaptic Systems for kindly providing m6A antibodies. This work was supported by a grant from the NIH to Y.S. (R01 GM117264), funds from Boston Children’s Hospital to Y.S. and Startup Funds provided by Harvard Medical School to K.A. Y.S. is an American Cancer Society Research Professor. This work used the Extreme Science and Engineering Discovery Environment (XSEDE), which is supported by NSF grant number ACI-1548562. Specifically, it used the Bridges system, which is supported by NSF award number ACI-1445606, at the Pittsburgh Supercomputing Center (PSC). D.V-G. is supported by a postdoctoral fellowship from the Mexican Council of Science and Technology (CONACYT, CVU: 257385) and is a member of the Mexican National System of Researchers (SNI I).

## Author contributions

E.S., D.V-G. and Y.S. conceived and designed the study. E.S., D.V-G., A.D., H.C., T.H. and W.S. performed the research. T.H. and K.A. performed and analyzed PRO-Seq. E.S., D.V-G. and Y.S. wrote the manuscript and all authors contributed to the final version of the manuscript.

## Data availability

Once the paper is published, all the genome-wide data will be accessible through GEO with accession number GSE122803.

## Methods

### Cell culture

MEL624 (RRID: CVCL_8054) and HEK293T (RRID:CVCL_0063) cells are grown in DMEM (Gibco) supplemented with 10% FBS and 1x pen/strep solution (Gibco) at 37°C in a CO_2_ incubator. Cells are split once they reach ∼90% confluence with a 1:8 ratio.

Cells are routinely checked for mycoplasma infection.

### mRNA purification

Total RNA is isolated from MEL624 cells with TRIzol reagent following manufacturer’s instructions. 200 μg of total RNA are subjected to two rounds of Poly(A) mRNA purification using the magnetic mRNA isolation kit from (NEB). 100 μL of oligo d(T) magnetic beads are used per purification. mRNA is eluted in a final volume of 50 μL, in elution buffer.

### Processing of RNA Samples for Mass Spectrometry

250 or 500 ng of mRNA is de-capped with a decapping enzyme in 20 μL reaction volume for an hour at 37°C. Then the RNA is digested with 0.5U of nuclease P1 (Sigma) in P1 buffer in 50 uL reaction at 37°°C for 2 hours. To dephosphorylate the single nucleotides, 1U of rSAP (NEB) is added to a final reaction volume of 100 μL with CutSmart buffer (NEB) and incubated for an hour at 37°°C. The 100 μL samples are filtered with Millex-GV 0.22u filters.

### HPLC-MS/MS Analysis

10 uL from each sample is injected into Agilent 6470 Triple Quad LC/MS instrument. The samples are run in mobile phase buffer A (water with 0.1% Formic Acid) and 2 to 20% gradient of buffer B (Methanol with 0.1% Formic Acid). MRM transitions are measured for adenosine (268.1 to 136.1, retention time 1.03 min), guanosine (284.1 to 152.1, retention time 1.15 min), 2’-O-methyladenosine (Am) (282.1 to 136.1, retention time 1.52 min), N6-methyladenosine (m6A) (282.1 to 150.1, retention time 1.79 min), N6,2-O-dimethyladenosine (m6Am) (296.1 to 150.1, retention time 2.40 min). The concentrations of each compound in the samples are calculated using calibration curves constructed with standard compounds of adenosine (Abcam), N6-methyladenosine (Abcam), N6,2’-O-dimethyladenosine (Toronto Research Chemicals). For LC/MS-MS data collection and analysis, Agilent Mass Hunter LC/MS Data Acquisition Version B.08.00 and Quantitative Analysis Version B.07.01 softwares are used.

### mRNA purification from different species

Total RNA was extracted with TRIzol from the following sources: Human: MEL624 melanoma cell line (RRID: CVCL_8054)

Mouse: C2C12 mice myoblast cell line (ATCC Cat# CRL-1772, RRID:CVCL_0188) Zebrafish: WT adult whole fish (ZFIN Cat# ZDB-GENO-990623-3, RRID:ZFIN_ZDB-GENO-990623-3)

Worm: Adult N2 (WT) *C. elegans* (WB Cat# N2, RRID:WB-STRAIN:N2) Fly: KC drosophila cell line (RRID:CVCL_Z833)

Yeast: *S. pombe*, ED666 h+ ade6-M210 ura4-D18 leu1-32 from Bioneer Inc. mRNA was extracted from 50-100 μg of total RNA as described above.

### Generation of PCIF1 KO cell lines

Two complementary strategies are followed to generate clonal PCIF1 MEL624 cell lines. 3 sgRNAs were cloned into the lentiCRISPR V2 vector (RRID:Addgene_52961):

PCIF1 sgRNA-1: TAGCGGTAAAGGAGCCACTG

PCIF1 sgRNA-2: CGGTTGAAAGACTCCCGTGG

PCIF1 sgRNA-3: ATTCACCAACCAGTCCCTGT

Additionally, a random non-targeting sgRNA was cloned into lentiCRISPR V2 as a control:

Random sgRNA-1: ATCGTTTCCGCTTAACGGCG

For the first strategy, 1 μg of lentiCRISPR V2 plus 0.5 μg of psPAX2 (RRID:Addgene_12260) and 0.2 μg of pMD2.G (RRID:Addgene_12259) packaging plasmids were mixed with 3 μg of polyethylenimine (PEI) in 100 ul of OptiMEM (Gibco) and added into 6-well plates of 30% confluent HEK293T cells (RRID:CVCL_0063). 4 hours post-transfection, the transfection media was replaced with 1.5 mL of regular media (DMEM, 10% FBS, 1X pen/strep). 48 hours post-transfection the media containing viral particles was passed through a 0.45 um filter and the viral particles from the 3 PCIF1 sgRNAs were combined. 200 μL of the pooled viral particles were used to infect a 6-well plate of ∼30% confluent MEL624. At the same time, another 6-well plate was infected with 200 μl from the non-targeting sgRNA. Under these conditions we reached an infection rate of ∼50%. 12 hours post-infection, cells were washed with PBS and media with 1 μg/mL of puromicyn was added to select infected cells. 48 hours post-selection, cells were grown with regular media for additional 48 hours and single-cell sorted in a 96-well plate in a FACS Aria cell sorter. Individual clones were validated as true KOs by western blot with a PCIF1 specific antibody (Bethyl A304-711A). This strategy generated PCIF1 KO1 and PCIF1 KO2 cell lines, as well as the control cell line from the non-targeting sgRNA clones.

In the second strategy, and to avoid integration of cas9 into the genome, 2 μg of lentiCRISPRv2 from the PCIF1 sgRNA-2 were transfected into a ∼30% confluent 6-well plate of MEL624 cells using 4 μl of lipofectamine 2000 according to manufacturer’s instructions. 24 hours post-transfection, cells were selected with 1 μg/mL of puromycin for 48 hours and growth for additional 48 hours without selection. Cells were single-cell sorted and evaluated as described above. This strategy generated the PCIF1 KO used for the rescues and m6Am mapping experiments.

### PCIF1 KO cell line rescue

WT and catalytic-deficient versions of PCIF1 coding sequence were cloned into the PHAGE-puro lentiviral plasmid with a C-terminal HA-FLAG tag. Lentiviruses were prepared as described above. 500 μL of lentiviral particles were used to infect ∼30% confluent MEL624 KO cells. 12 hours post-infection, viruses were washed out with PBS and cells were selected with media containing 1 μg/mL of puromycin for 48 hours. After selection, rescue cell lines were continuously growth in media containing 0.5 μg/mL of puromycin to avoid the silencing of PCIF1 transgene.

### PCIF1 immunofluorescence

PCIF1 KO and rescue cell lines were seeded in a 96-well plate to a ∼70% confluence. Cells are fixed for 10 mins in 3.7% formaldehyde in PBS at room temperature. Cells are washed twice with PBS and permeabilized for 10 mins with 0.1% saponin in PBS. Cells are blocked at room temperature for 1 hour with IF blocking buffer (10% BSA, 0.3% triton in PBS) and then are incubated overnight at 4°C with a monoclonal anti-FLAG antibody (Sigma-Aldrich Cat #F1804, RRID:AB_262044) diluted 1:500 in IF buffer (10% FBS, 1% BSA, 0.3% triton in PBS). Primary antibody is washed three times with 0.05% PBS-Tween. Cells are incubated with Alexa 488 goat anti-Mouse (Thermo Fisher Scientific Cat #A-11029, RRID:AB_2534088) 1:1000 in IF buffer for 1 h at room temperature in the dark. DAPI is added to a final concentration of 0.2 μg/mL and incubated for additional 10 mins. Cells are washed three times with 0.05% PBS-Tween, once with PBS and then imaged in a Nikon Eclipse Ti fluorescent microscope. Pictures of representative fields are taken with a S Plan Fluor 20X/0.45 objective with the NIS Elements v4.20 software (DAPI channel exposure: 500 msec. GFP channel exposure: 600 msec.).

### Thin Layer Chromatography

1 μg of mRNA is cleaned, labeled and digested as previously described (Kruse et al., 2011; Mauer et al., 2017). 10% of the starting material is loaded into a TLC cellulose glass plate and first and second dimensions are run for 14 and 19 hours, respectively. Plates are exposed to a phosphor imager screen for 24 hours and read in a Typhoon scanner. Pictures are analyzed and quantified using Fiji v2.0.0 (RRID:SCR_002285).

### Recombinant Protein Purification

Full-length human PCIF1 gene is cloned into pGEX-4T1 vector. It is expressed overnight in 500 mL Rosetta bacteria (Novagen) culture induced with 0.1 mM IPTG at 18°C. The cells are lysed in 15 mL cold lysis buffer (50 mM Tris pH 7.4, 150 mM NaCl, 0.05% NP-40, 1mM PMSF) with 0.25 mg/mL chicken lysozyme on ice for 30 min. The cell lysate is sonicated on ice and cleared with centrifugation for 20 min at 10000 rpm at 4°C. The GST tagged recombinant protein is bound to Glutathione Sepharose 4B beads (GE Healthcare) at 4°C with rotation for 3 hours. The beads are washed with lysis buffer containing 500 mM NaCl. The untagged recombinant protein is eluted off the beads with overnight rotation at room temperature with 5U of thrombin in thrombin cleavage buffer (50 mM Tris pH 8.0, 150 mM NaCl, 5mM CaCl_2_). The protein preparation is supplemented with 10% glycerol and stored at −80°C.

Full-length human EIF4E gene with N terminal 6xHis tag is cloned into pGEX-4T1 vector. The same purification protocol for recombinant PCIF1 is followed with the following exceptions. The initial lysis buffer is supplemented with 10U RNaseA/T1 mix. The cleared lysate is first bound to Ni-NTA beads at 4°C for 3 hours to purify using the 6xHis tag, washed in lysis buffer with 750 mM NaCl and eluted in His elution buffer (50 mM Tris pH 7.4, 150 mM NaCl, 150 mM imidazole). The eluted protein is bound to Glutathione Sepharose 4B beads (GE Healthcare) at 4°C with rotation for 1 hour and eluted off the beads with overnight rotation at room temperature with 5U of thrombin in thrombin cleavage buffer (50 mM Tris pH 8.0, 150 mM NaCl, 5mM CaCl_2_). The protein preparation is supplemented with 10% glycerol and stored at −80°C.

### Enzyme Kinetics

The enzymatic parameters for m7G-Am methylation were determined by incubating full-length untagged recombinant PCIF1 enzyme (3 nM) with increasing concentrations of RNA dinucleotide (10 – 500 nM) at 37 C in 50 mM Tris-HCl (pH 7.5 @ 25 °C), 5 mM β-ME, 10 mM EDTA and 80 μM S-adenosylmethionine (SAM) buffer. Aliquots were withdrawn at t = 0, 2, 5, 10, 30 and 60 min and boiled for 3 min to stop all enzymatic activity. The aliquots were further processed as described in Processing of RNA samples for mass spectrometry methods section. The levels of the final product (m7G-m6AmG) formed at each time point was determined using mass spectrometry by monitoring the amount of m6Am in each aliquot. The observed rate of product formation (kobs) was determined by plotting the concentration of m6Am against time for each concentration of the substrate (10, 25, 50, 100, 250, 500 nM). The kobs vs substrate concentration curve was fit to the Michaelis-Menten equation using the Graphpad Prism software to obtain the final enzymatic parameters. The same protocol was followed for kinetics using EGFP mRNA with indicated concentrations.

### *In vivo* translation assays

200 ng of either Am-, or m6Am-EGPF mRNA are transfected into a 6-well plate with MEL624 at ∼30% confluence. Transfection is done with 1 μl of lipofectamin 2000 according to manufacturer’s instructions. Untransfected wells are used as negative controls. 24 h post-transfections, cells are imaged in a Nikon Eclipse Ti fluorescent microscope and pictures of representative fields are taken with a S Plan Fluor 20X/0.45 objective with the NIS Elements v4.20 software (GFP channel low exposure: 500 msec. GFP channel high exposure: 3 sec. bright field exposure: 20-40 msec.). At the same time, a second set of transfected wells is trypsinized, washed once with PBS and resuspended in 500 μl of PBS for flow cytometry. Cells are analyzed in a FACS Canto II and percentages of GFP positive cells as well as GFP intensity values are calculated with FloJo 8.7. Finally, a third set of transfected wells is lysed with 200 μL of TRIzol and total RNA is purified following manufacturer’s instructions. Quantitative PCR is performed with primers specific for GFP and GAPDH as internal control in order to determine the level of mRNA that was transfected into the cells.

### *In vitro* Translation

In vitro translation reactions are carried out using Retic Lysate IVT kit (Invitrogen Ambion) in 25 μL or 10 μL reactions with varying indicated mRNA concentrations (50, 100 or 200 ng). The translation is carried out at 30°C for 1 hour. For the firefly luciferase mRNA translation reactions, luciferase reagent supplied by Firefly Luc One-Step Glow Assay Kit (Pierce Thermo Fisher Scientific) is used. For the in vitro transcribed bicistronic firefly-IRES-renilla luciferase mRNA translation reactions, luciferase reagents supplied by Dual-Luciferase Reporter Assay System (Promega) are used and luminescence is measured on a plate reader.

### m6A MeRIP-Seq

m6A MeRIP experiments were performed following the recommendations from the EpiMark N6-Methyladenosine Enrichment Kit (NEB #E1610). Briefly: 10 μg of mRNA were incubated at 98°C for 3 min in fragmenting buffer (10 mM Tris pH 7.4, 10 mM ZnCl_2_) to generate fragments of ∼100-150 bp. Fragmented RNA is incubated with protein G magnetic beads conjugated to 1 μL (250 μg) of anti-m6A rabbit monoclonal antibody from the EpiMark kit for 1 h at 4°C. Beads were washed twice with 200 μL of reaction buffer, twice with 200 μL of low salt buffer and twice with 200 μL of high salt buffer. Immunoprecipitated RNA was eluted with TRIzol and purified. Illumina sequencing libraries are prepared with the NEBNext Ultra II Directional Library Prep Kit for Illumina from NEB (#E7760), following manufacture’s instructions. RNA-Seq libraries are generated from three biological replicates for Input (RNA-Seq) and m6A-immunoprecipitated mRNA. Libraries are sequenced in a HiSeq 2000 with an average coverage of 20 million reads per sample.

Reads are aligned against hg38 genome assembly with hisat 2.0.4 using the following parameters: --no-unal --rna-strandness R. SAM files are sorted and converted to BAM with samtools v1.7. Reads with QS < 10 are excluded, Normalized bigWig files are generated from BAM files with bamCoverage from the deepTools suite v3.0.2 with the following parameters: -bs 20 --normalizeUsing BPM --skipNAs. Reads that overlap with the blacklisted regions from ENCODE project are excluded. Average bigWig files for the biological replicates are generated using the bigwigCompare program from the deepTools suite v3.0.2. m6A-enriched genes are obtained by calling significant peaks form the BAM files with macs2 (v2.1.1) with a q-value < 1e-10 for each individual biological replicate. A gene is considered to be enriched for m6A only if a peak was called in all the replicates. Metagene and heatmap plots were generated with the deepTools suite v3.0.2 and in-house scripts.

### PRO-Seq sequencing and analysis

PRO-Seq libraries from 3 independent biological replicates were generated for Control MEL624 cells, PCIF1-KO clone 1 and PCIF1-KO clone 2, using 1 million cells per sample. Samples were spiked with 40,000 Drosophila S2 cells as a normalization spike-in control. Samples were sequenced on the NextSeq using a High Output 75-cycle kit, generating paired end 42 nt reads. Libraries were sequenced to an average read depth of 50 million mappable reads per sample.

Paired-end reads were trimmed to 40 nt. To remove adapter sequence and low quality 3’ ends we used cutadapt 1.14, discarding reads shorter than 20 nt (-m 20 -q 10), and removing a single nucleotide from the 3’ end of all trimmed reads to allow successful alignment with Bowtie 1.2.2. Remaining read pairs were paired-end aligned to the *Drosophila* dm3 genome index to determine spike-normalization ratios based on uniquely mapped reads. Counts of pairs mapping uniquely to spike-in RNAs (drosophila genome) were determined for each sample. In this case, the samples displayed highly comparable recovery of spike-in reads, thus only depth normalization was used for each bedGraph.

Reads mapped to dm3 were excluded from further analysis, and unmapped pairs were aligned to the hg38 genome assembly. Identical parameters were utilized in each alignment described above: up to 2 mismatches, maximum fragment length of 1000 nt, and uniquely mappable, and unmappable pairs routed to separate output files (-m1, -v2, -X1000, --un). Pairs mapping uniquely to hg38, representing biotin-labeled RNAs were separated, and strand-specific counts of the 3’-end mapping positions determined at single nucleotide resolution, genome-wide, and expressed in bedGraph format with “plus” and “minus” strand labels, as appropriate. Combined bedGraphs were generated after deduplication by summing counts per nucleotide of all 3 replicates for each condition.

Read counts were calculated per gene (from transcription start site to transcription end site), in a strand-specific manner, based on default annotations (ensembl hg38), using featureCounts. Differentially expressed genes were identified using DESeq2 v1.18.1 under R 3.3.1. PRO-Seq size factors were determined based on DESeq2 (for Control: 0.9690449, 0.8752877, 0.9008983; PCIF1-KO1: 1.1669854, 0.9802904, 1.3105898; PCIF1-KO2: 0.9866223, 1.0047620, 0.9516695). At an adjusted p-value threshold of <0.01, 731 affected genes in KO clone 1 and 1406 in KO clone 2 were identified as differentially expressed upon PCIF1-KO in MEL624 cells. After overlapping both lists there are 506 genes that pass the padj < 0.01 threshold, from which 320 are upregulated in both KO clone 1 and clone 2 (Table S4), 182 are downregulated in both clones (Table S5) and 4 are downregulated in KO1 and upregulated in KO2. UCSC Genome Browser tracks were generated from the combined replicates per condition, normalized as in the differential expression analysis.

### RNA-Seq analysis

Read counts for the RNAseq inputs from the m6A mapping experiment are calculated from BAM files with featureCounts from the Rsubread package v1.22.3 in an R 3.3.3 environment. The read count table is analyzed with edgeR v3.12.1 and log2 TPM measurements are generated. After calculating normalization factors and estimating dispersion, DEG genes between KOs and Control cells were calculated using glm modeling with a FDR < 1e-5. A final list of DEG shared between PCIF1 KO1 and PCIF1 KO2 is generated using in-house scripts, which gives a total of 236 upregulated genes (Table S2) and 305 downregulated genes (Table S3). Boxplots of average expression are generated with R 3.3.3 and in-house scripts from the log2 TPM values calculated in edgeR. Mann-Whitney tests to compare average expression levels between different samples and hyper geometric test to calculate p-values for overlaps between datasets in Venn diagrams are performed in R 3.3.3.

### Microscale Thermophoresis (MST)

RNA oligos used in MST analysis correspond to the first 219 nt 5’ end of bicistronic mRNAs used in Figure 7F. The 219 nucleotide m7G-Am capped RNA oligos are transcribed using a PCR generated DNA template using the bicistronic construct plasmid (pFR_CrPV_xb). Am methylated RNA oligo is further in vitro methylated to m6Am using recombinant PCIF1. To generate decapped RNA control, m7G-Am methylated RNA is decapped using a decapping enzyme. The RNA constructs are column purified using Zymo RNA Clean & Concentrator kit after each transcription, in vitro methylation step and decapping reaction.

The full-length human recombinant N terminal 6xHis tagged EIF4E protein is fluorescently labelled in PBST with 0.1% PEG8000 using His-Tag Labelling Kit RED-Tris-NTA (2nd Generation) (Nanotemper MO-L018). The RNA oligos are denatured at 65°C for 3 minutes followed by 1 minute immediate ice incubation. Then 2-fold serial dilutions of RNA are carried out in PBST with 0.1% PEG800 at room temperature using LoBind microfuge tubes. Labelled protein is added at 25 nM dye and 100 nM protein final concentration to each RNA dilution. After 5 minute room temperature incubation, premium capillaries are loaded and MST is carried out with 100% LED power and 40% laser (20 seconds ON and 5 seconds OFF) using NanoTemper Control 1.1.9 software. The binding constants are calculated using MO.Affinity Analysis V2.3 software using 10 dilution points carried out in triplicate independent binding assays for each RNA oligo.

## Supplemental Information

**Figure S1.**
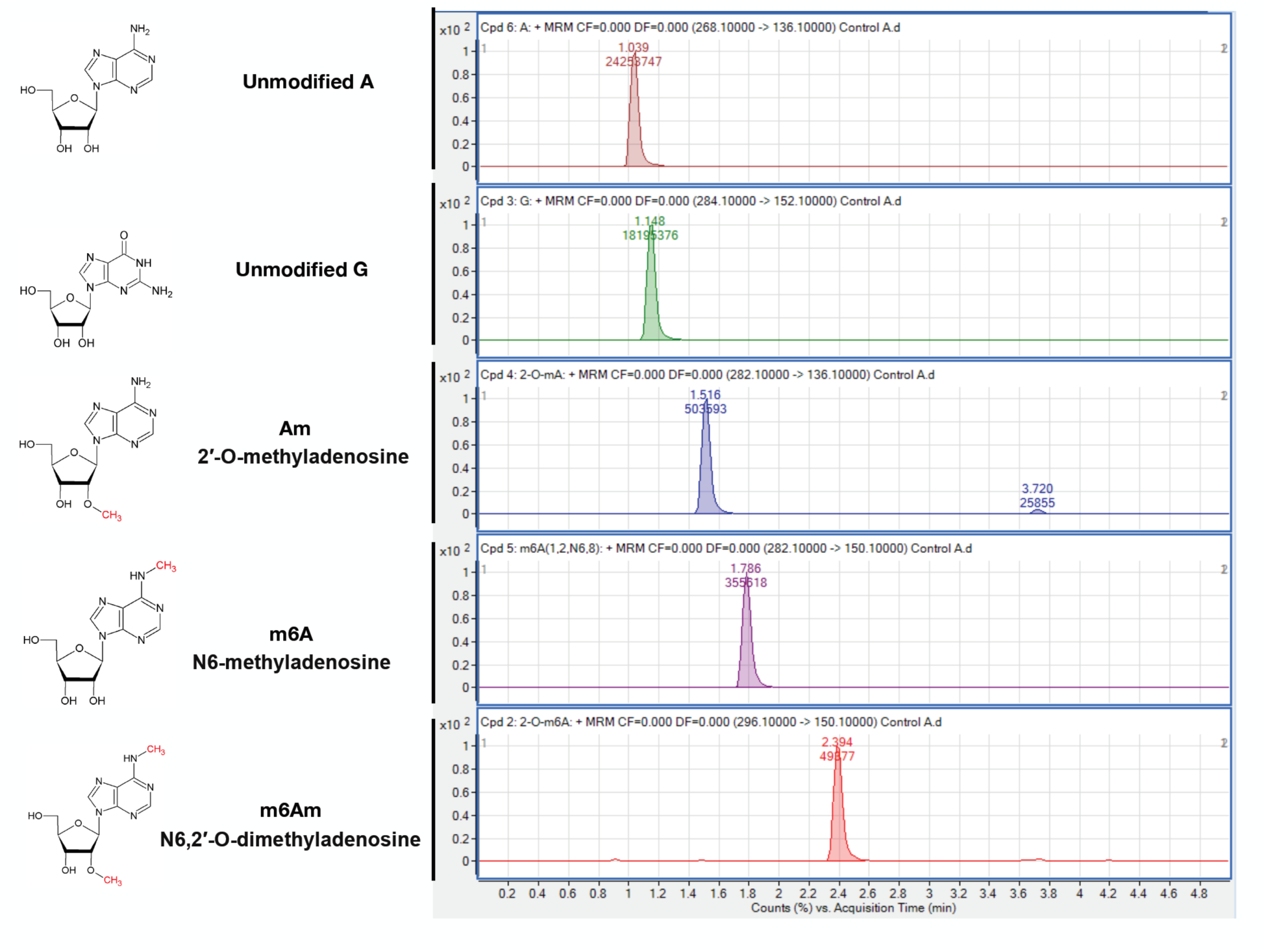
Related to Figure 1. Representative MS spectra of de-capped mRNA from Mel624 human melanoma cell line.

**Figure S2.**
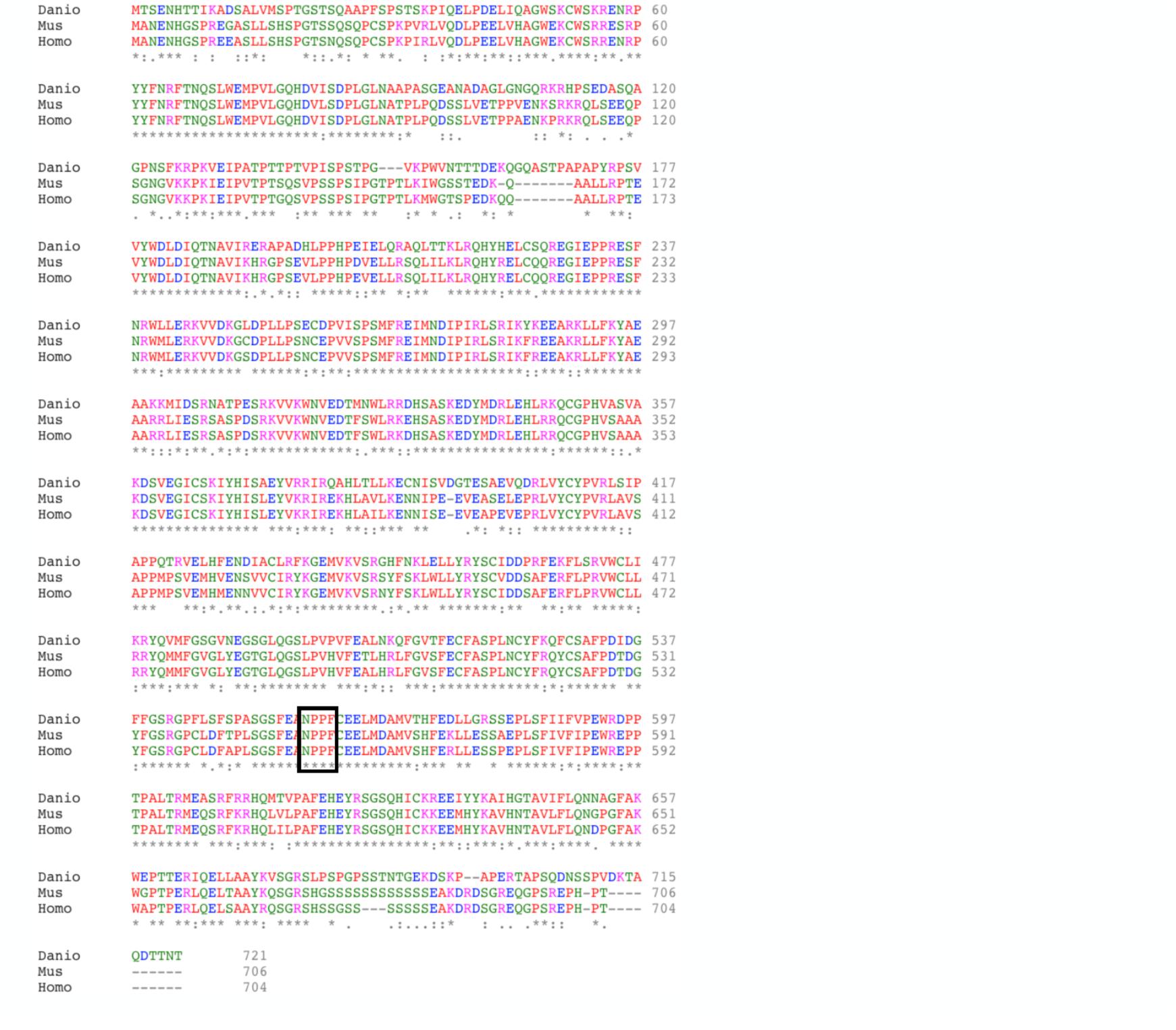
Related to Figure 2. Alignment of protein sequences of zebrafish, mouse and human PCIF1 using ClustalW. The conserved methyltransferase motif is highlighted.

**Figure S3.**
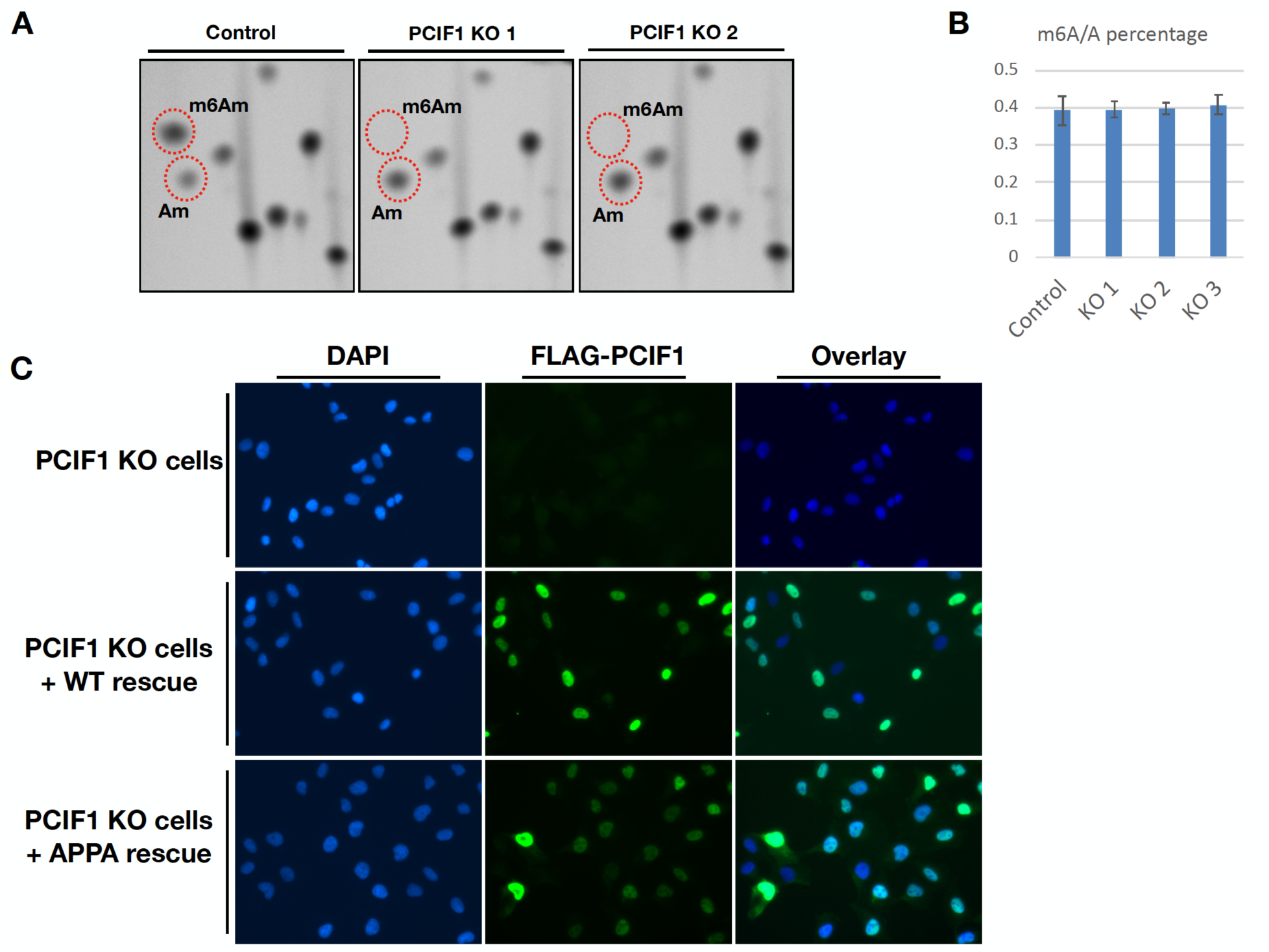
Related to Figure 2. PCIF1 is required for mRNA m6Am methylation in vivo. 3A. Thin-layer chromatography analysis of cap-adjacent nucleotides from mRNA isolated from control and two independent PCIF1 KO cell lines. 3B. LC-MS/MS m6A/A percentages of mRNAs from control and three independent PCIF1 KO cell lines

3C. Immunofluorescence images of PCIF1 KO and FLAG-PCIF1 rescue cell lines with anti-FLAG antibody.

**Figure S4.**
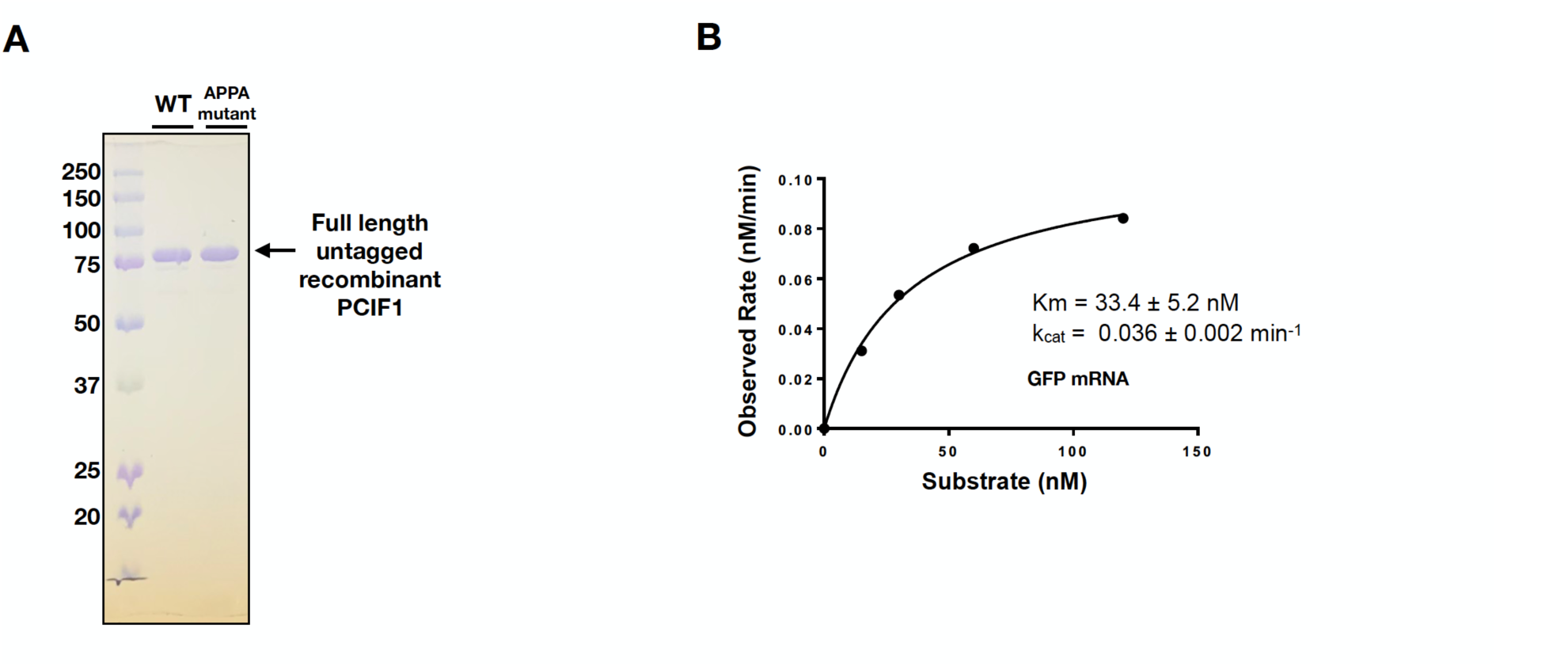
Related to Figure 3. *In vitro* enzyme kinetics with recombinant PCIF1. 4A. GST tagged PCIF1 was expressed in bacteria and purified using Glutathine Sepharose beads and eluted using overnight thrombin cleavage. Around 3 μg of purified recombinant PCIF1 was run on PAGE gel and stained with Coomassie Blue. 4B. Michaelis–Menten kinetics of recombinant PCIF1 is determined using m7G-Am capped EGFP mRNA.

**Figure S5.**
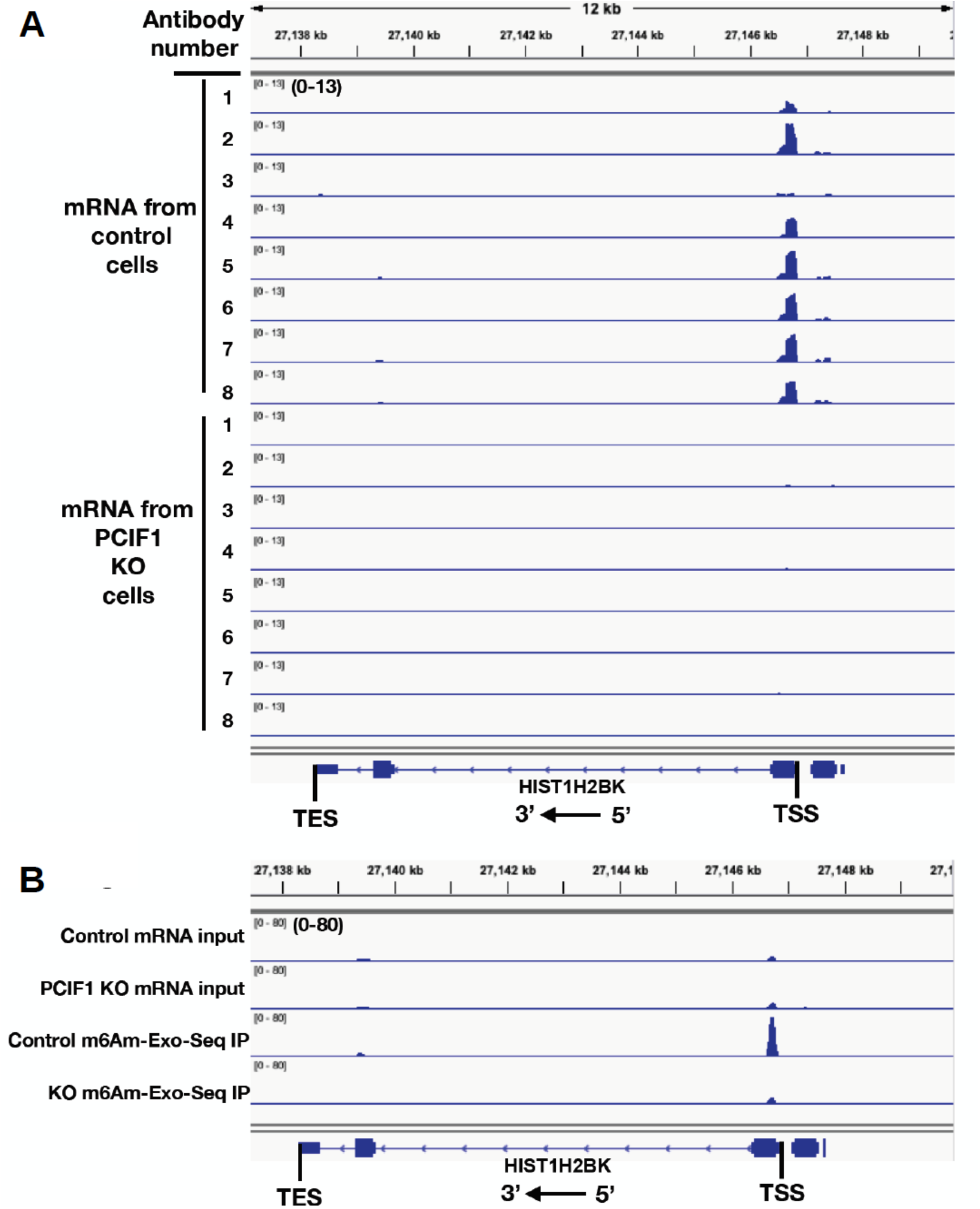
Related to Figure 5. Global mapping of m6Am through m6Am-Exo-Seq. 5A. Genome browser view of m6Am-Exo-Seq fold change over input employing 8 different antibodies against m6A in control and PCIF1 KO cells. 5B. Genome browser view of reads per million in inputs and corresponding m6Am-Exo-Seq IPs from control and PCIF1 KO cells.

**Figure S6.**
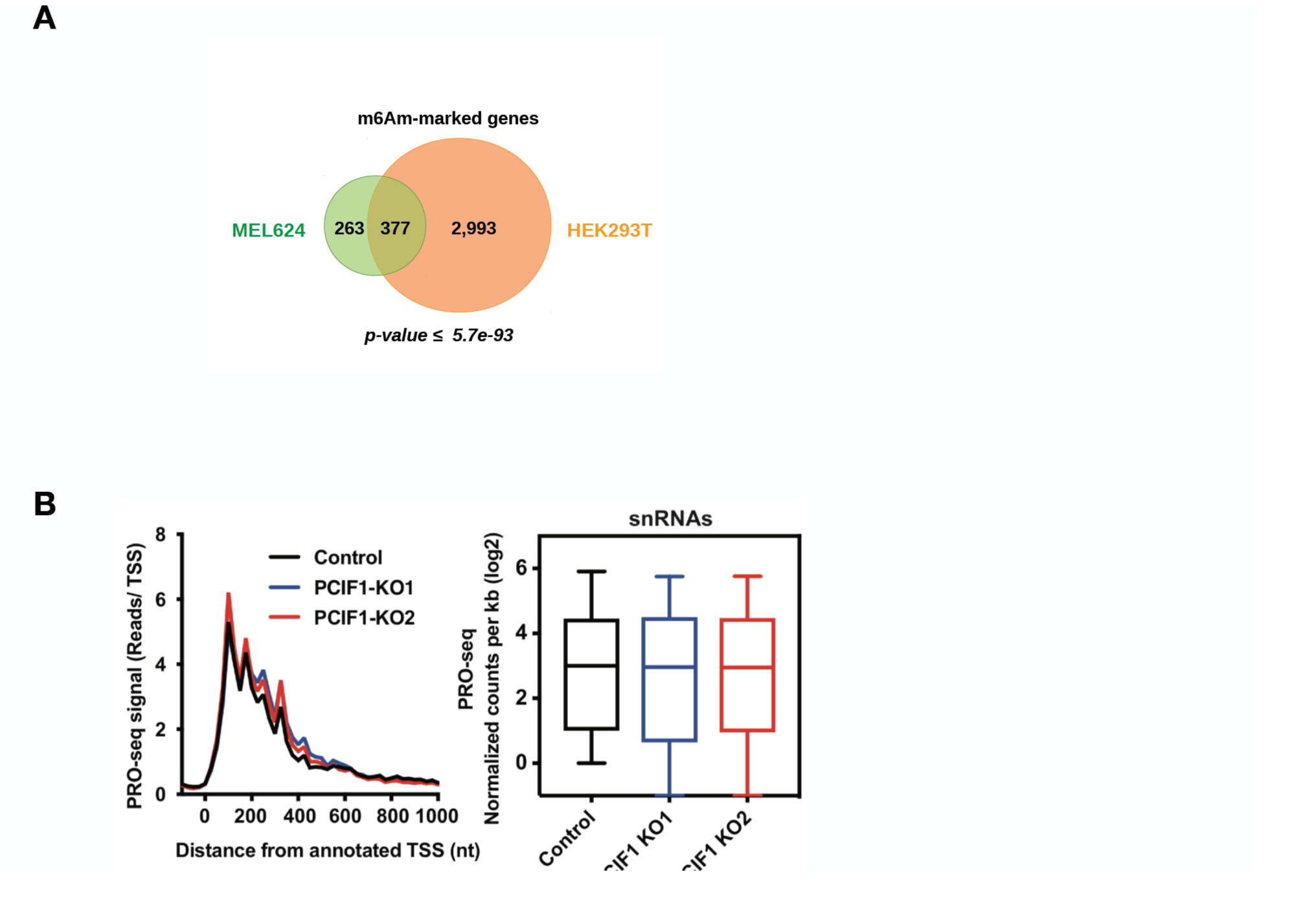
Related to Figure 6. m6Am is cell type specific and does not affect snRNA transcription. 6A. Venn diagram depicting overlap between 640 m6Am enriched genes in MEL624 and 3370 identified in HEK293T cells. P-values are calculated with a hypergeometric test. 6B. PRO-Seq signal is shown at snRNA genes (N=667 snRNA genes expressed in MEL624) in control and PCIF1 KO cells. Shown are a composite metagene distribution (left) or boxplot of PRO-Seq signal within snRNA genes. Boxplots are shown as in 7D. No significant differences were detected.

**Figure S7.**
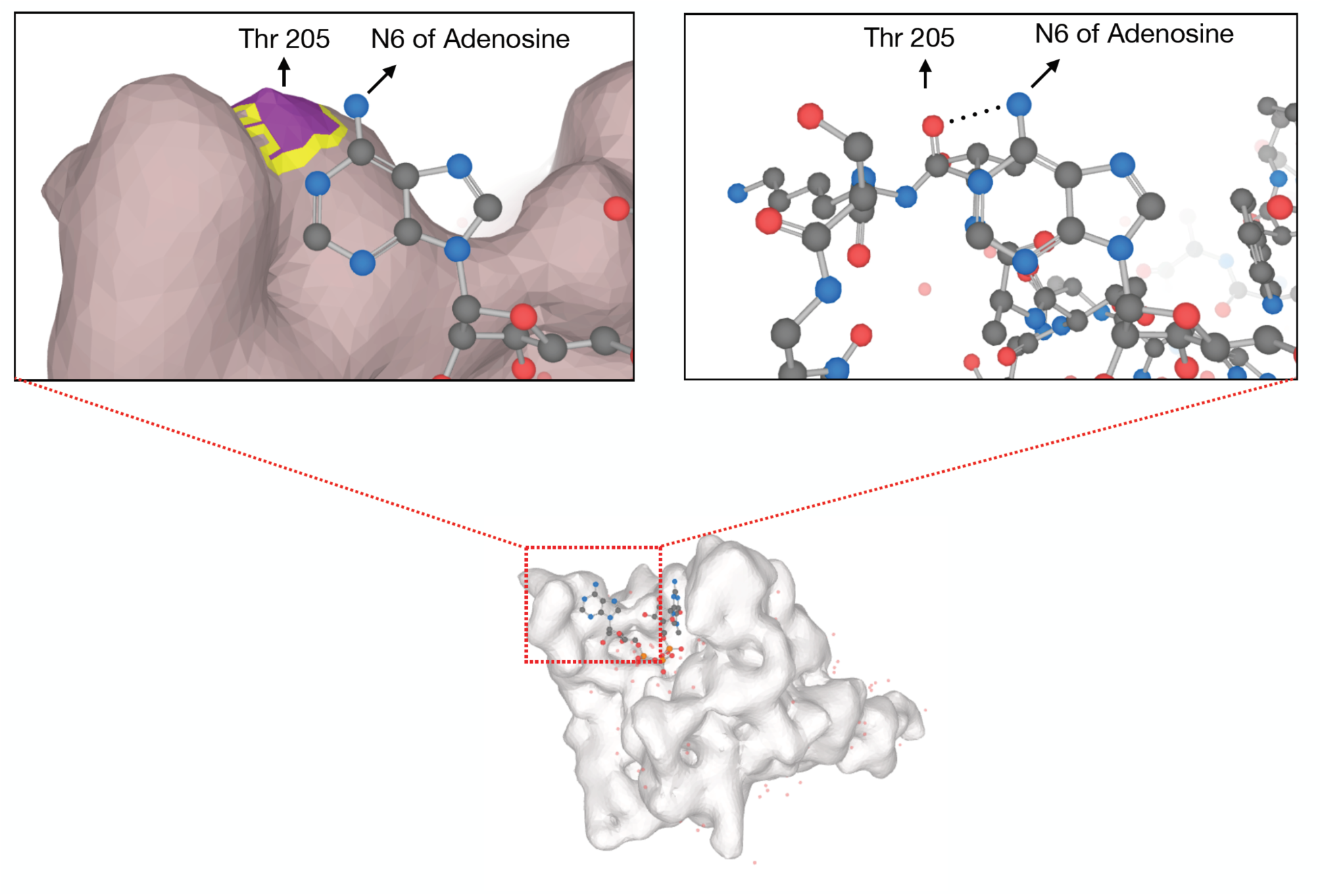
Related to Figure 7. N6 residue of cap adjacent adenosine of mRNA, which gets m6A methylated by PCIF1, forms a hydrogen bond with threonine 205 residue of human eIF4E.

**Figure S8.**
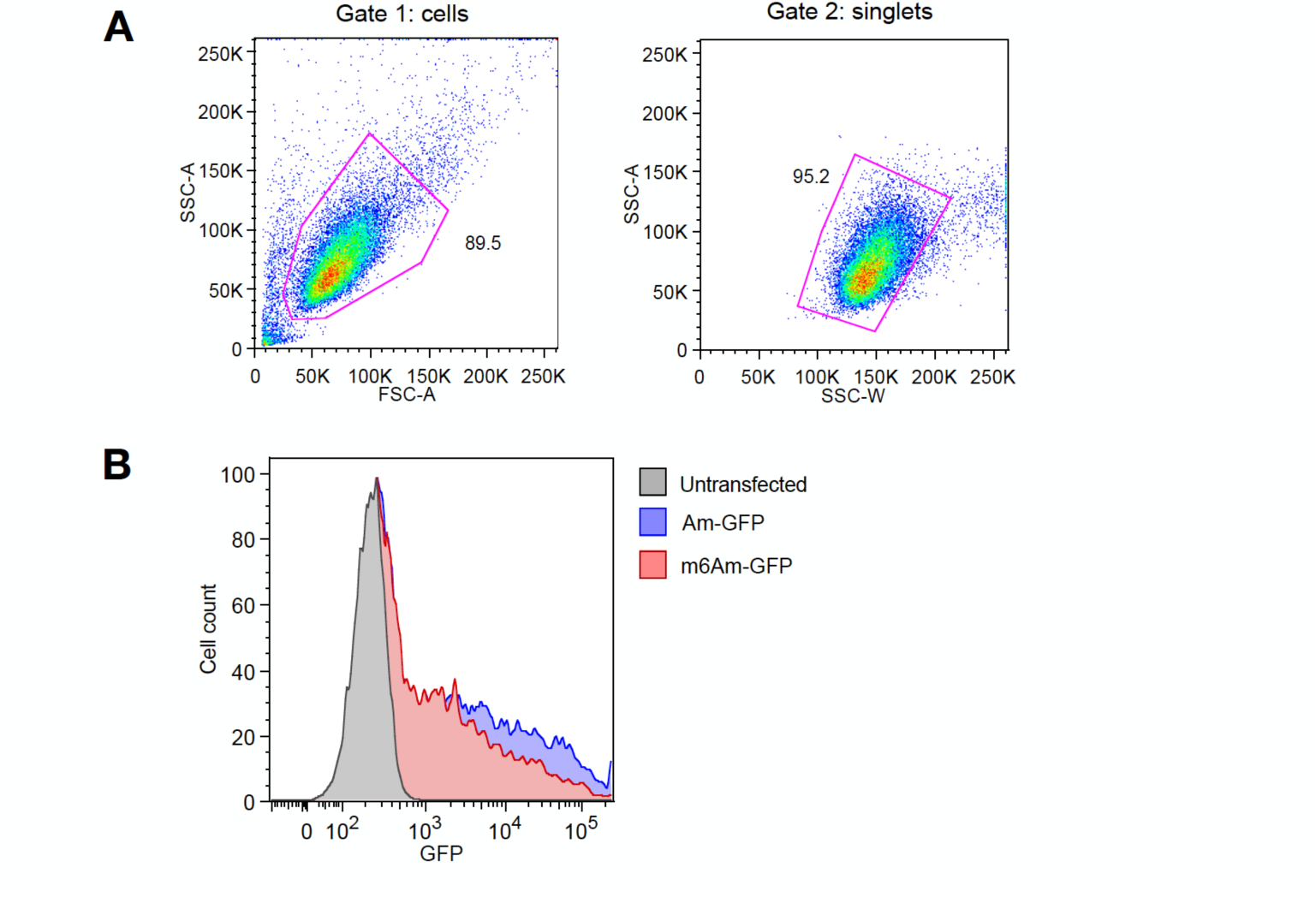
Related to Figure 7. FACS analysis of Am or m6Am EGFP transfected MEL624 cell lines. 8A. Gating strategy for selecting singlet cells. Cells are gated based on FCS-A and SSC-A (gate 1). Singlets are selected from gate 1 by gating SSC-W vs SSC-A. Cells from gate 2 are used for all quantifications shown in Figure 7. 8B. Representative fluorescent profile of MEL624 cells transfected with Am- or m6Am-EGFP mRNA. Untransfected cells are used as negative control.

**Figure S9.**
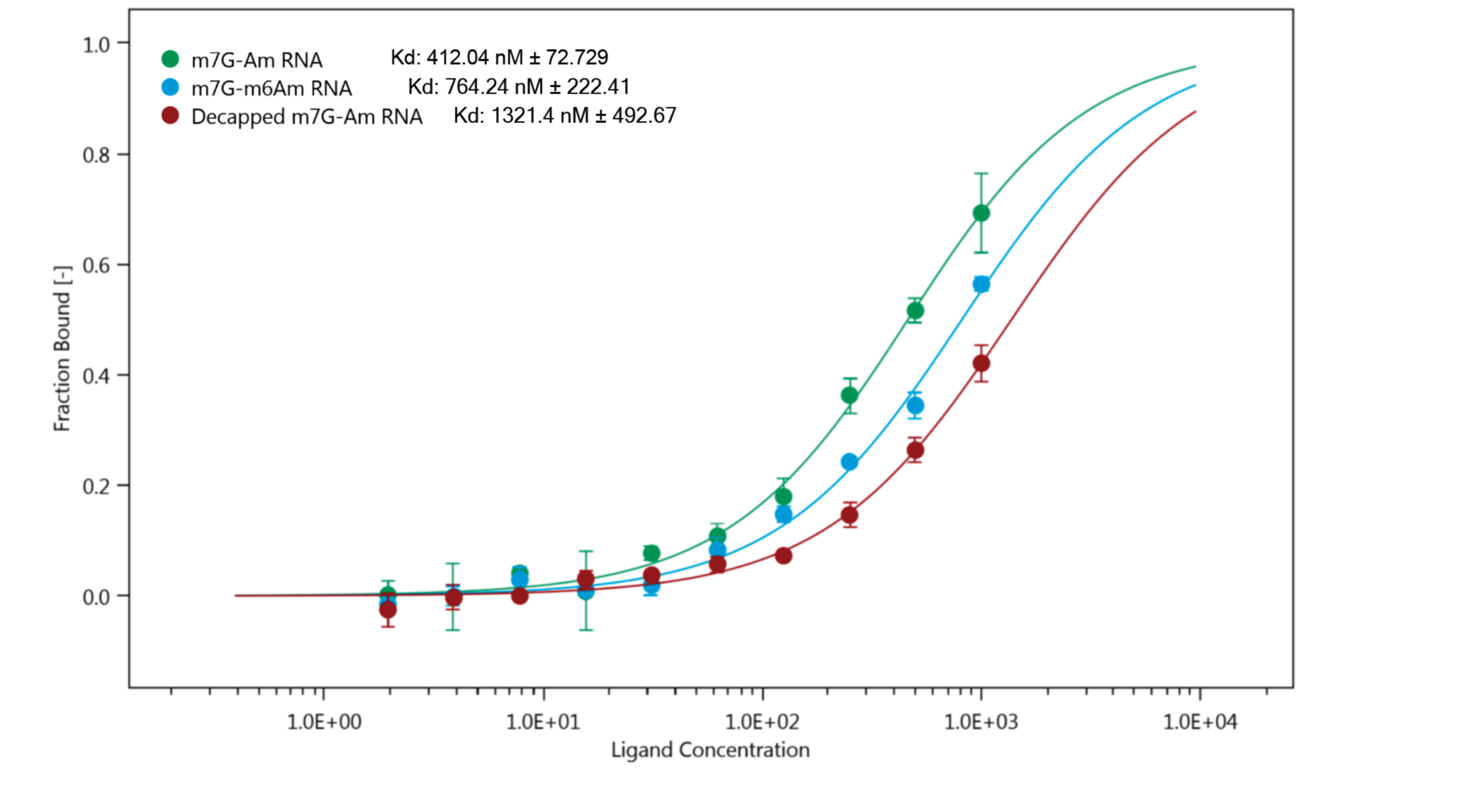
Related to Figure 7. Kd constants of fluorescently labelled N-terminal His tagged recombinant human eIF4E with 217 nt RNA oligos with either m7G-Am or m7G-m6Am 5’ ends measured by microscale thermophoresis. De-capped m7G-Am RNA was used as a control. The 217 nucleotide RNA oligo sequence corresponds to the first 217 nt of bicistronic construct that was used in in vitro translation assays in Fig.7F. Three independent binding assays were performed with freshly labelled protein for each RNA oligo.

**Table S1. Related to Figure 5.** List of m6Am-marked genes.

**Table S2. Related to Figure 6.** List of Upregulated genes according to RNA-Seq

**Table S3. Related to Figure 6.** List of Downregulated genes according to RNA-Seq

**Table S4. Related to Figure 6.** List of Upregulated genes according to PRO-Seq

**Table S5. Related to Figure 6.** List of Downregulated genes according to PRO-Seq

